# The transcriptional landscape of glycosylation-related genes in cancer

**DOI:** 10.1101/2022.09.28.509853

**Authors:** Ernesto Rodriguez, Dimitri Lindijer, Sandra J. van Vliet, Juan J. Garcia Vallejo, Yvette van Kooyk

## Abstract

Changes in glycosylation patterns have been associated with malignant transformation and clinical outcome in several types of cancer, although no comprehensive analysis has been performed in a pan-cancer setting. Here, we performed an extensive transcriptomic analysis of glycosylation related genes (such as enzymes involved in synthesis and degradation of glycoconjugates, transporters, mucins and galectins), using publicly available bulk and single cell transcriptomic data sets from tumor samples and cancer cell lines. We identified genes and pathways associated with different tumor types, which may represent novel diagnostic biomarkers as α2-3 sialylation for Melanoma, MUC21 for Lung adenocarcinoma and Galectin-7 for Squamous cell carcinomas (SCC). Accordingly, serum levels of Galectin-7 in patients with lung cancer were elevated in SCC respect to adenocarcinomas, supporting its biomarker potential. Moreover, we characterized the contribution of different cell types to the overall glycosylation profiles observed by performing the integration and analysis of 14 single cell RNA-seq datasets. This led us to identify that cancer cells are responsible for the specific tumor glyco-codes identified in bulk transcriptomics, while stromal and immune cells contribute in a conserved manner across various malignancies. Furthermore, our results suggest that the glycosylation-related genes and pathways expressed by cancer cells are influenced by the cell of origin and the oncogenic pathways that led to malignant transformation. Lastly, we described the association of different glycosylation-related genes and pathways with the clinical outcome of patients. Interestingly, while the expression of genes associated to some pathways (as proteoglycan biosynthesis) are consistently associated with a more aggressive disease, the correlation of others pathways with the survival of patients depends on the particular tumor type. Remarkably, the expression of genes associated with the *synthesis of CMP-sialic acid* was correlated with lower survival of patients in Uveal Melanoma and PDAC, while the opposite was observed for colorectal cancer. The extensive transcriptomic analysis of glycosylation pathways in cancer that we report here can serve as a resource for future research aimed to unravel the glyco-code in cancer related to clinical outcome or biomarker development.

## Introduction

Glycosylation is a complex and highly regulated metabolic pathway that leads to the enzymatic synthesis of glycan structures attached to different macromolecules such as proteins, lipids or RNA, collectively known as glyco-conjugates^1-3^. In the human genome, hundreds of genes encode for proteins that directly or indirectly contribute to the diverse array of glycan structures found in the cell (also denominated *glycome*)^4^. Alterations in the glycosylation signatures are a universal feature of malignant transformation and different cancer-associated glycan structures have been associated with tumor progression, by playing key roles in cell-to-cell adhesion, modulation of signaling receptors, metastasis and induction of the tolerogenic microenvironment^2,3,5^. Moreover, several biomarkers used in the clinic for diagnostic or prognostic purposes are glycan structures or glycoproteins, as is the case of CA19-9, a carbohydrate structure also known as sialyl Lewis a (sLe^a^), which serum levels are increased in gastrointestinal cancers^3,6.^

One of the most studied glycan carriers in cancer are the mucins, proteins containing hundreds of glycosylation sites in its backbone. In normal tissue, mucins are expressed in the apical membrane of polarized epithelial cells; however, cancer cells often lose their polarization, resulting in the presence of mucins in the tumor microenvironment and the blood stream^5^. In fact, several mucins are currently used as biomarkers in the clinic, as MUC1 (CA15-3) in breast cancer and MUC16 (CA125) in ovarian cancer^3,7,8^.

Additionally, glycan structures can be recognized by carbohydrate binding proteins, also called lectins. In humans, almost two hundred genes encode for lectins that differ in their glycan specificity and cell type expression^2,4.^ Galectins are a family of soluble lectins with specificity towards glyco-conjugates containing β-galactose that can be secreted by tumor and stromal cells, functioning as autocrine or paracrine signals that work in a glycan-dependent manner^9^. They have been functionally involved in a broad range of biological processes in cancer, as RNA splicing, anti-tumor immune response and cell migration and differentiation, among others^9^. In the last decade, the extensive use of transcriptomic analysis in cancer research has revolutionized our current understanding of the disease, generating an important amount of publicly available datasets^10,11.^ However, the role of glycosylation has been widely overlooked. Here, we characterized the expression of glycosylation-related genes (GRGs) and pathways (GRPs) in a pan-cancer setting using bulk and single cell transcriptomic datasets from tumor tissues and cell lines. This allowed us to identify GRGs and GRPs associated with different tumor types, characterize the differential contribution of cancer and stromal cells to the overall tumor glyco-code and study their association to the clinical outcome of patients.

## Results

### Identification of cancer-specific glycosylation signatures

To perform a glycosylation focused transcriptomic analysis of cancer, we started by selecting the glycosylation-related genes (GRGs). For this, we performed a literature search to pinpoint the genes encoding for <u>*enzymes*</u> involved in the synthesis or degradation of glyco-conjugates and activated sugar donors, as well as the ones corresponding to glycan-specific membrane <u>*transporters*</u> (Figure 1A, Supplementary Table 1). We also included genes that encode for the protein backbone of <u>*mucins*</u>, carriers of cancer associated glycan-structures that can be present in circulation; as well as <u>*galectins*</u>, as they can function as glycan-dependent signaling molecules in the tumor microenvironment (Supplementary Table 1). Moreover, we made use of gene sets (both custom-made and from different databases) for the transcriptomic analysis of glycosylation-related pathways (GRPs) using gene set enrichment analysis (Supplementary Table 2).

**Figure 1.**
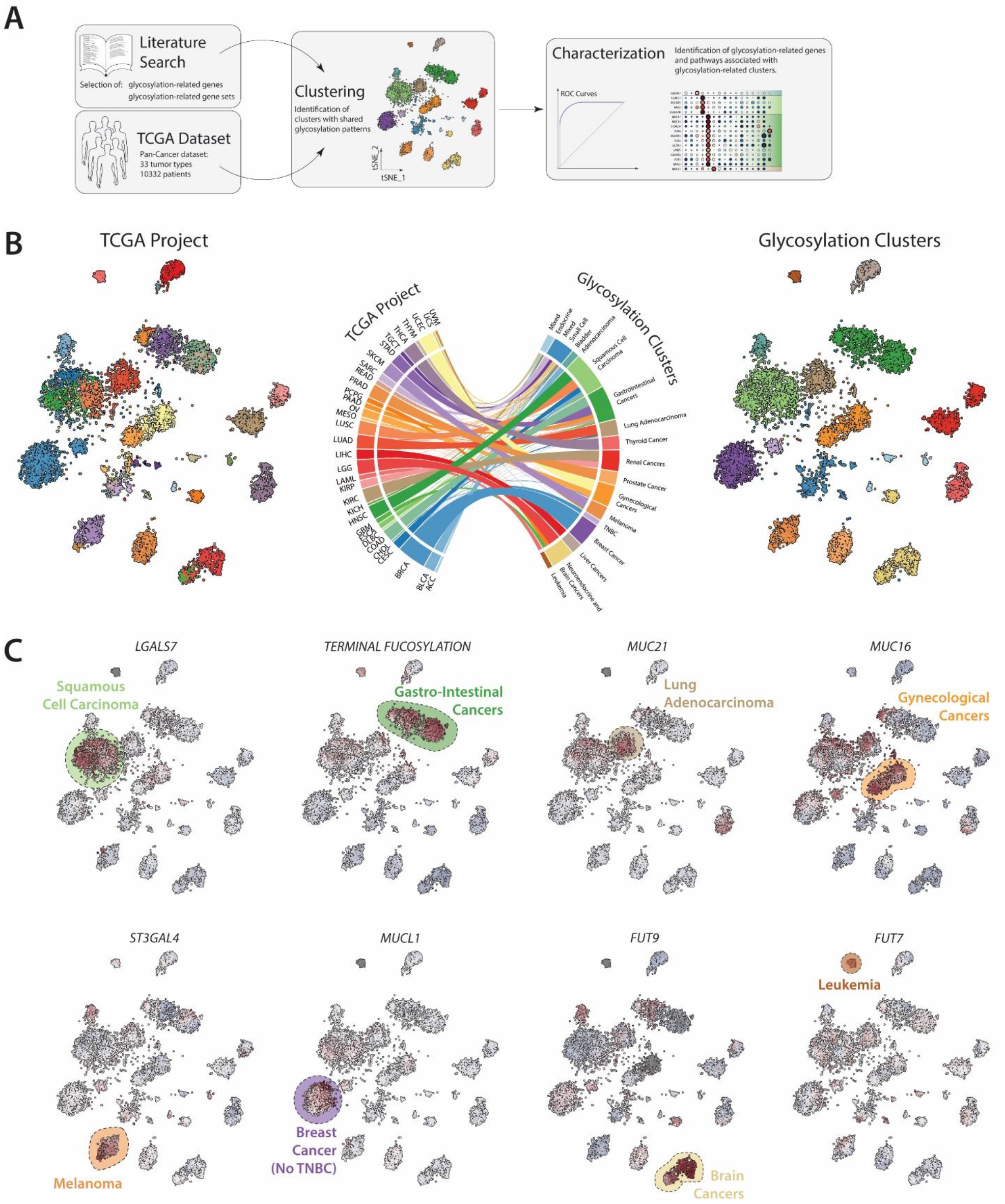
Identification of specific glycosylation signatures of tumor. A) Workflow used in this paper for the transcriptomic analysis of glycosylation-related genes in cancer. B) Consensus clustering of the TCGA pan cancer data set using glycosylation-related genes. TSNE plot showing TCGA projects (left) and glycosylation-clusters (rights). Cord diagram showing the association of the glycosylation clusters with the TCGA project (center). C) Expression of selected specific glycosylation-related genes in the different clusters.

Firstly, we explored we whether we could identify GRGs and GRPs to be associated with different cancer types. For this, we used the TCGA pan cancer dataset which contain bulk RNA-Seq data from 10322 samples derived from 33 different tissues (Supplementary Table 3). Given that transcriptomics signatures can be shared between samples from different tumor types, we started our analysis by performing consensus clustering using the glycosylation-related genes. This analysis led to the identification of 16 clusters of tumor samples with similar GRG expression (Figure 1B). In line with previous analysis, the tissue of origin heavily influences the clustering, with samples of a given tissue generally grouped together in the same glycosylation cluster, with the exception of a series of heterogeneous cancers, such as Breast (BRCA), Bladder (BLCA), Cervical (CESC), and Esophagus (ESCA) cancers (Supplementary Figure 1)^10^. This is also reflected in the clusters that present only samples from the same organ or tissue type, as is the case for Thyroid (THCA), Renal (renal clear cell [KIRC] and papillary cell [KIRP] carcinomas), Prostate (PRAD) and Blood (acute myeloid Leukemia [LAML]) cancers (Figure 1B, Supplementary Figure 1). This suggests that the main determinant of GRG pattern in tumors is dictated by the tissue in which the tumor originates.

We also observed samples that are grouped based on morphological features or tissue/organ similarity: S*quamous Cell Carcinomas* (SCC; light green), are grouped together independent of the tissue in question (Lung, Head and neck, Cervix, Esophagus or Bladder); same for *Gastro-Intestinal cancers* (colon [COAD], rectal [READ], stomach [STAD] and pancreatic [PAAD] cancers; green); *Gynecological cancers* (ovarian [OV] and endometrial [UCEC] cancers; orange); *Melanomas* (skin cutaneous [SKCM] and uveal [UVM], dark orange), derived from both the eye and the skin; *Liver cancers*, including cholangiocarcinomas (LIHC and CHOL, respectively; gray); *Neuroendocrine and Brain Cancers* (pheochromocytoma and paraganglioma [PCPG], low grade gliomas [LGG] and glioblastoma [GBM]; yellow). Interestingly, we also observed that samples of patients with *Triple Negative* Breast cancer (TNBC; light purple) group together to form an independent cluster from the rest of breast cancer subtypes (Figure 1B). We also observed two mixed clusters, which we denominate as *Endocrine* (adrenocortical carcinoma [ACC], chromophobe renal cell carcinoma [KICH] and thymoma [THYM]; light blue) and *Small Cell* (B-cell lymphoma [DLBC], mesothelioma [MESO], sarcomas [SARC] and testicular germ cell tumors [TGCT]; blue).

To identify signatures associated to each cluster, we used ROC curves and calculated prediction power based on the area under the curve (Figure 1C and Supplementary Figure 2 and 3). This led to the identification of genes and pathways that were specific for each cluster: *MUC16* (CA125), specifically expressed in *Gynecological cancers*, and “TERMINAL_FUCOSYLATION”, specific for *Gastro-Intestinal cancers*, which includes enzymes involved in the synthesis of Lewis antigens, as sialyl-Lewis^a^ (sLe^a^, CA19-9). This suggest that some of the novel signatures identified in this analysis could also serve as potential biomarkers for the different cancer types, for example: Galectin-7 (encoded by the genes *LGALS7, LGALS7B*) for Squamous cell carcinomas; *MUC21* for Lung Adenocarcinomas, *ST3GAL4* for Melanomas, *MUCL1* for Breast cancer (No TNBC), *FUT9* for Brain cancers and *FUT7* for Leukemia (Figure 1C).

To explore the use of the GRGs and GRPs expression signatures as potential biomarkers, we shifted our focus to the group of cancers arising from the same tissue that ended up in different clusters based on the expression of GRGs (Figure 2A, Supplementary Figure 1). With the exception of Breast cancer (which is separated TNBC and non-TNBC), the samples in these tissues are grouped based on their histological subtypes, separating SCC (all of which are clustered together) from other subtypes, as adenocarcinomas. In Esophagus carcinoma (ESCA), adenocarcinoma samples cluster with the *Gastro-Intestinal cancers*, while in Lung and Cervical cancer, Adenocarcinomas are grouped together (Figure 2A). In Bladder cancer (BLCA), the *Luminal* molecular subtype forms its own cluster and the *Neural* is associated with the *Mixed – Small Cell* (Figure 2A). Given that SCC and adenocarcinomas are associated with differences in the clinical outcome of patients (Figure 2B), we next investigated if differences in the expression of GRGs and GRPs can be used as a biomarker to distinguish these tumor types. In particular, we focused in galectin-7 for the differential diagnosis of non-small cell lung cancers (NSCLC) given its highly upregulation in SCC, but not adenocarcinoma, when compared to normal tissue (Figure 2C). In a small cohort of NSCLC patients (containing 16 SCC and 20 adenocarcinomas), we measured the plasma levels of galectin-7 using ELISA and observed a higher concentration in SCC than in Adenocarcinoma (Figure 2D). Thus, our data suggest that galectin-7 is a good diagnostic biomarker candidate for SCCs, supporting the idea that this analysis can serve as a resource for potential discovery of new biomarkers.

**Figure 2.**
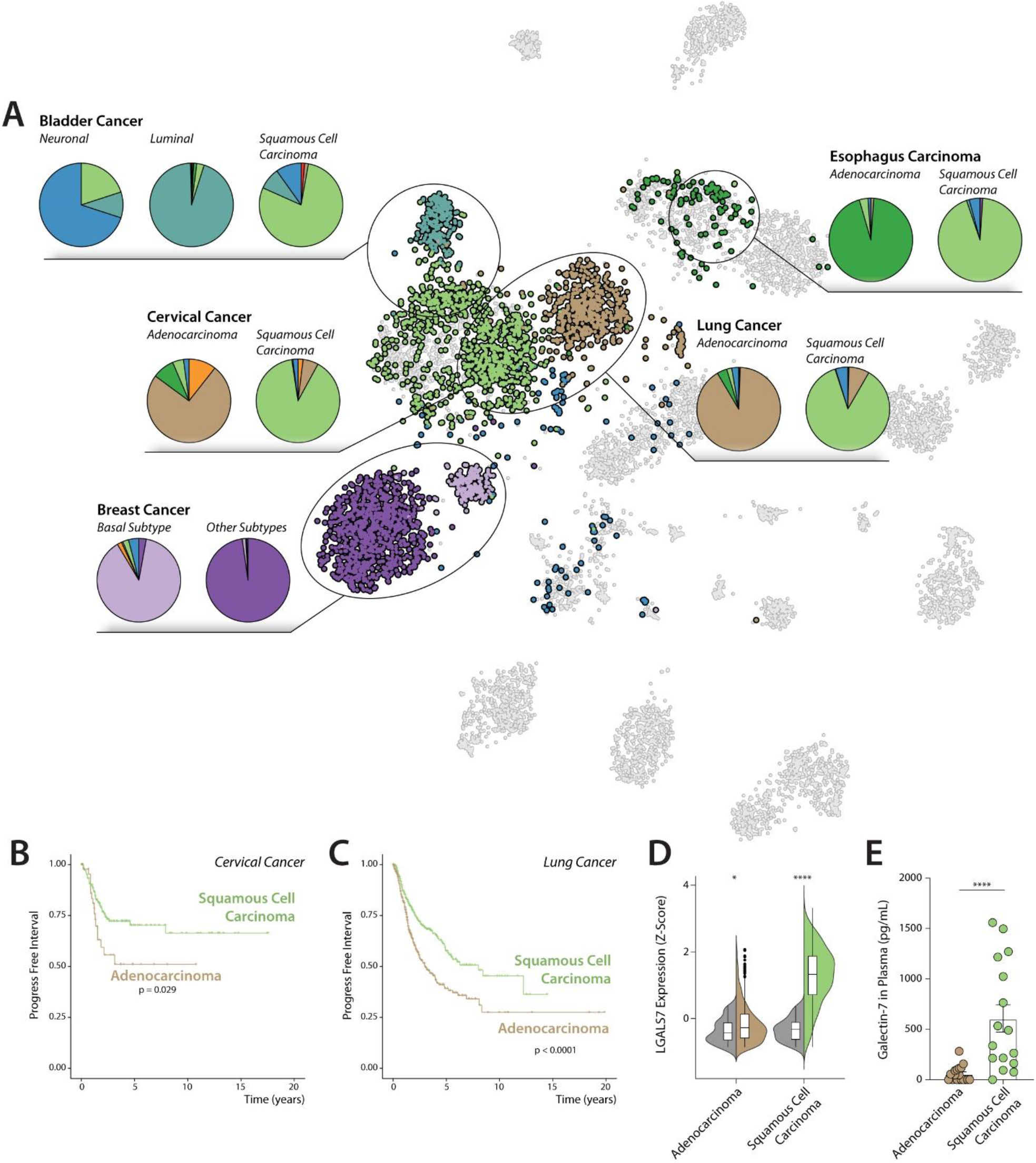
Glycosylation signatures distinguish Adenocarcinomas and Squamous cell carcinomas in different tumor types. A) Association of glycosylation-related cluster with different diagnosis or molecular subtypes in heterogenous cancers. Progression free survival between Adenocarcinoma and Squamous Cell Carcinoma in cervical cancer (B) and lung cancer (C). D) Expression of LGALS7 in normal and tumor tissue in Adenocarcinoma (n_normal_ = 59, n_tumor_ = 510) and Squamous Cell Carcinoma (n_normal_ = 51, n_tumor_ = 502) of the lung. E) Plasma concentration of Galectin-7 in Adenocarcinoma (n = 20) and Squamous Cell Carcinoma (n = 16) of the lung. In violin plot: data presented as boxplot indicate the median, 25th and 75th percentiles (hinges) and whiskers represent 1.5 times the interquantile range.

### Cancer cells are the main contributors to the tumor glycosylation signature

The limitation of analyzing bulk transcriptomic data is the lack of cellular resolution for determining the relative contribution of the different cell populations to the tumor glycosylation patterns. To overcome this limitation, we analyzed single cell RNA-Seq (scRNA-Seq) from 14 different tumor types, representing 9 different TCGA glycosylation clusters and containing data from 250 patients (Figure 3A-B)^12-26^. These datasets were individually pre-processed, down sampled to 10.000 cells, and integrated into a single dataset using the Reciprocal principal component analysis (RCPA) algorithm as implemented in the package *Seurat*^27^. Dimensionality reduction and clustering of this integrated data set led to the identification of common populations, as immune cells (T and B lymphocytes, Myeloid and Mast cells), other stromal cells (as Fibroblasts, Pericytes and Endothelial cells) and a cluster that contained malignant cells and some tissue specific cells (Figure 3C, Supplementary Figure 4). This cluster was selected and individually cleaned up for each dataset to discard tissue specific cells and wrongly clustered cells. To analyze the relative contribution of the cancer and stromal compartment to the overall expression of glycosylation-related genes in each tumor type, we performed differential gene expression between malignant cells and all the other cell types present in tissue (Figure 3D, Supplementary Figure 5). This analysis led to the identification of genes generally associated with cancer cells (*MUC1, LGALS3, UGDH, TSTA3* and *GALE*) or the stromal compartment (*LGALS1, PSAP, LGALS9, IDS, MGAT1*) in most of the tumor types analyzed (Supplementary Figure 5). Interestingly, we observed that despite the strong association of Galectin-1 (LGALS1) and *PSAP* with the stromal compartment, cancer cells appear to be the main source expressing these genes in skin cutaneous melanoma (Supplementary Figure 5). We also found GRGs that are associated with cancer cells in a tissue specific manner, including some of the genes that were identified in our previous analysis in bulk transcriptomics (Figure 3D, Supplementary Figure 5). For example, galectin-7 genes (*LGALS7* and *LGALS7B*) are expressed by cancer cells in SCC; fucosylation-related genes (*GMDS, FUT2, FUT3*) in Gastro-Intestinal cancers; *MUC21* in Lung Adenocarcinoma; *MUC16* in ovarian cancer; *ST3GAL4* and *ST3GAL6* in cutaneous and uveal Melanoma; *GALNT6* and *MUCL1* for non-TNBC breast cancer and *ST6GAL1* and *UGT2B4* for Liver Cancer (Figure 3D). These data show that cancer cells are the main contributors to the specific tumor glycosylation signatures found in tissue.

**Figure 3.**
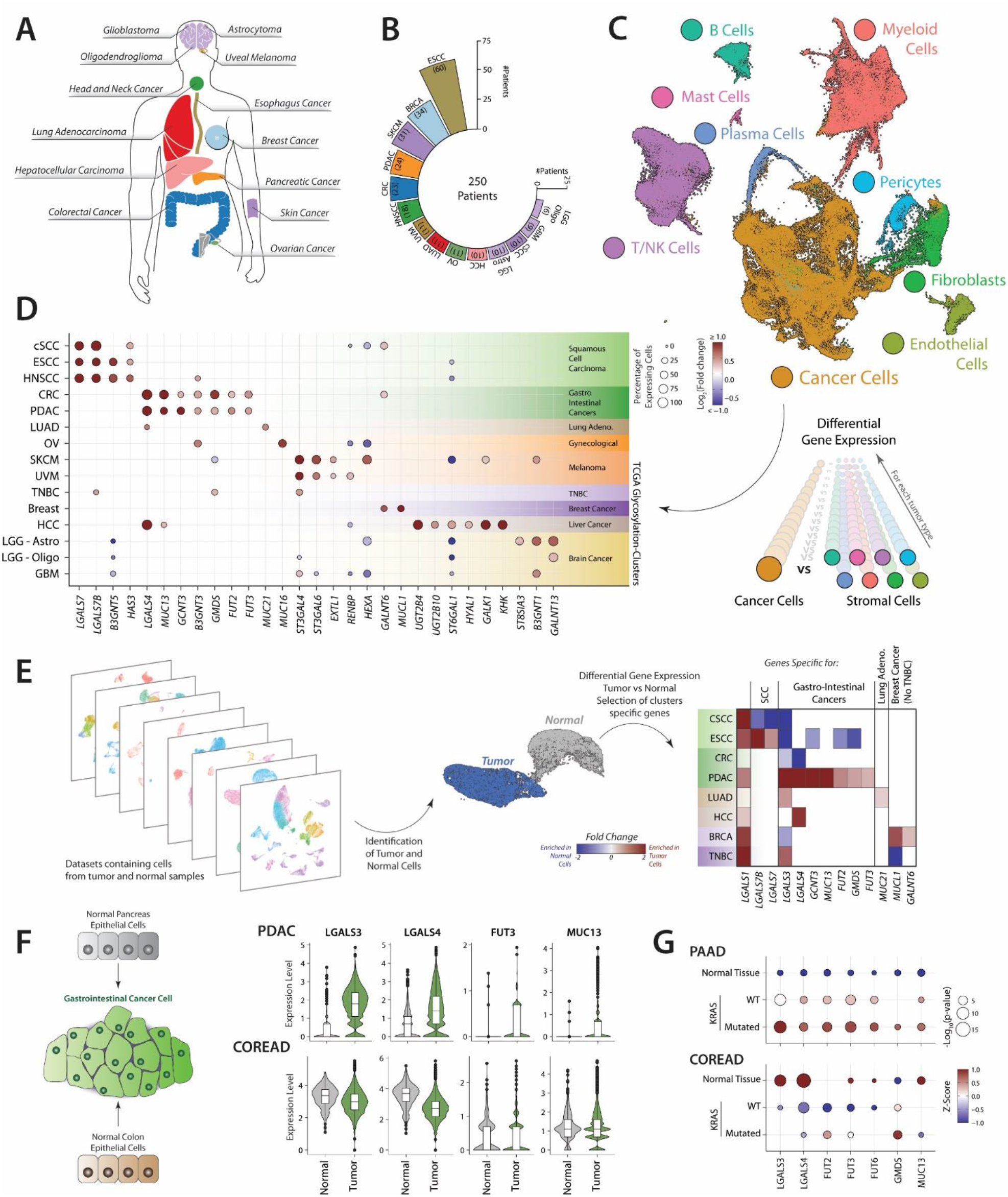
Cancer cells are main contributors to the specific tumor GRG expression signatures and is dependent on the cell of origin. A) Tumor types analyzed by scRNA-Seq. B) Number of patients analyzed in the different tumor types. C) UMAP plot of the integrated scRNA-Seq. D) Dot Plot of selected differentially expressed GRGs between Cancer and stromal cells in the integrated scRNA-Seq. E) Differential expression analysis of GRGs between normal and cancer cells in individual scRNA-Seq datasets. F) Violin plot showing the expression of selected GRGs in Normal and Tumor cells from Pancreatic (PDAC) and Colorectal (COREAD) cancer. G) Dot plot displaying the differences between Normal and cancer tissue with or without mutations in KRAS. Statistical differences were evaluated using the Kruskal-Wallis test of one condition against the others.

The strong association of TCGA glycosylation-clusters with tumor type and the main contribution of cancer cells to their specific expression patterns, raises the question whether the cell of origin is a major determinant of cancer glycosylation signatures. To study this, we analyzed the expression of specific glycosylation-related genes in normal and cancer cells from tumors whose cell of origin has been identified. In cutaneous SCC, that originates from keratinocytes in the skin, we observed expression of galectin-7 genes (*LGALS7* and *LGALS7B*) in both normal and cancer cells (Supplementary Figure 6A). The same is the case for epithelial cells from normal colon or colorectal cancer, which express Galectin-3 (*LGALS3*) and 4 (*LGALS4*); hepatocytes from normal tissue and hepatocellular carcinoma (HCC) express *ST6GAL1* and *UGT2B4*; and melanocytes derived from cutaneous SCC (healthy) and cutaneous Melanoma (malignant), that both show expression of *ST3GAL4* and *ST3GAL6* (Supplementary Figure 6A). These data suggests that the cell of origin influences the GRG expression pattern of cancer cells and highlights the importance of studying changes in expression of glycosylation-genes between normal and cancer cells.

We next performed a differential gene expression analysis between cancer and normal cells in single cell RNA-Seq data containing samples from healthy and tumor tissue (Figure 3E, Supplementary Figure 6B)^13,18,20,23,26^ This allowed us to identify changes in GRG expression patterns specific for cancer cells, without being influenced by the different cell distributions that can be present in bulk transcriptomics. The gene *LGALS1* was found as the most commonly upregulated transcript in cancer cells of different tumor types (Figure 3E). We also found that some GRGs specifically associated with different tumor types are upregulated when compared to normal cells, as is the case of *MUC21* in Lung adenocarcinoma and *GALNT6* and *MUCL1* in non-TNBC Breast cancer. However, for some GRGs, the changes observed in the expression of GRGs depend on the tissue of origin. That is the case for genes encoding for galectin-7 (*LGALS7* and *LGALS7B*), highly expressed in SCC, that were found to be downregulated in cancer cells from cutaneous SCC but upregulated in the ones from esophagus SCC (Figure 3E). The same is the case for Galectin-3 (*LGALS3*) and Galectin-4 (*LGALS4*), associated with gastro-intestinal cancers, which are downregulated in colorectal (COREAD) but upregulated in pancreatic cancer (PAAD). Other GRGs highly expressed in gastro-intestinal cancers (as *GCNT3, GMDS, FUT2, FUT3* and *MUC13*) are only found differentially upregulated in PAAD but not in COREAD (Figure 3E). This shows that despite that gastro-intestinal cancers present similar patterns of expression of GRGs, their expression dynamic from normal cells nevertheless depends on the cell of origin (Figure 3F). We hypothesized that the presence of shared oncogenic pathways in these tumor types drive malignant cells to express a similar profile of GRGs independent of the level observed in the cell of origin. Given that PAAD and COREAD are characterized by the presence of KRAS mutations, we next analyzed if its presence in tissue leads to a similar expression profile as observed in scRNA-Seq^28^. Our data revealed that the expression of gastro-intestinal cancer GRGs are upregulated in PAAD samples with KRAS mutations, but not in COREAD, in line with our hypothesis (Figure 3G, Supplementary Figure 7).

### The contribution of the stromal compartment to the GRGs expression profile

Despite that cancer cells are the main source of the specific GRGs and GRPs found in different tumor types, our results also show that the stromal compartment can contribute glycosylation profile present in the tissue (Supplementary Figure 5). To confirm if what is observed in the scRNA-Seq analysis is also reflected in different tumor types present in the TCGA dataset, we correlated the expression of GRGs with the stromal fraction as described previously (Figure 4A)^29^. We confirmed that the stroma-associated genes identified in the previous analysis (as *LGALS1, PSAP, LGALS9* and *IDS*; Supplementary Figure 5) were also positively correlated with the stroma fraction in bulk RNA-Seq (Supplementary Figure 8A). However, in this new analysis we were able to identify other stroma-associated GRGs, such as *FUT7, HK3, LGALS2* and *CHST11* (Figure 4A). If we plot the expression of some genes in the integrated scRNA-Seq dataset we can observe different patterns of expression, with *MNFG* and *ST8SIA4* found in immune cells and *LGALS9* and *LGALS2* mainly restricted to Myeloid cells (Figure 4B-C). These results suggest that different stromal cells present in tumor tissue have particular expression profiles of GRGs.

**Figure 4.**
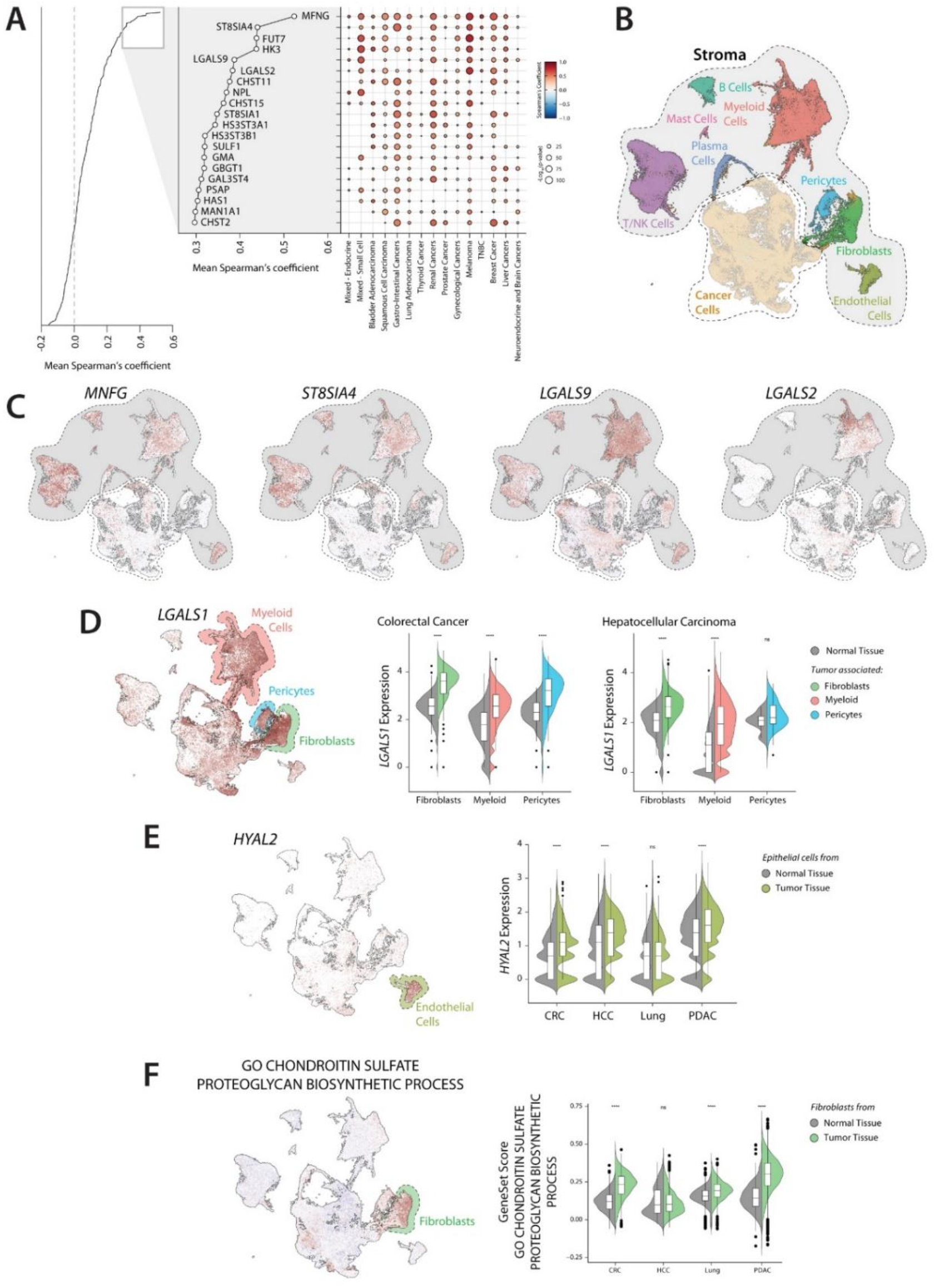
Stromal contribution to the tumor glycosylation. A) Correlation of glycosylation-related genes with the stromal fraction in the TCGA data set. B) UMAP plot of the integrated scRNA-Seq data set displaying the stromal cells. C) Expression of the stromal associated genes *MNFG, ST8SIA4, LGALS9* and *LGALS2* in the scRNA-Seq data. D) Expression of *LGALS1* in the scRNA-Seq data. Violin plot illustrating the expression of *LGALS1* in Fibroblasts, Myeloid and Pericytes from normal or tumor tissue of Colorectal cancer and Hepatocellular carcinoma. E) Expression of *HYAL2* in Epithelial cells from normal or tumor tissue in different tumor types. F) Expression of the gene set “*Chondroitin sulfate - Proteoglycan biosynthetic process*” in Fibroblasts from normal or tumor tissue in different tumor types. Statistics: Wilcoxon test (*p ≤ 0.05; **p ≤ 0.01; ***p ≤ 0.001; ****p ≤ 0.0001). In violin plot: data presented as boxplot indicate the median, 25th and 75th percentiles (hinges) and whiskers represent 1.5 times the interquantile range.

We next characterized the GRGs expressed by the different stromal cells using a modified version of the function *FindConservedMarkers* of the package *Seurat* to perform differentially gene expression in the integrated scRNA-Seq dataset (Supplementary Figure 8B-C). This allowed us to identify the specific expression of *CHPF, UGDH* and *HEXB* in Fibroblasts; *HYAL2, GALNT18* and *SULF2* in Endothelial cells; *PSAP, LGALS9, MGAT1* and *NPL* in Myeloid cells; *MGAT4A* in T/NK Cells; *GALC* in Mast Cells and *ST6GAL1* in B and Plasma cells (Supplementary Figure 8C). We also observed GRGs that are shared by different cell types, such as *SUFL1, LGALS3BP, UCGC* and *ST6GALNAC6* in Pericytes and Fibroblasts or *IDS* in B and T Lymphocytes (Supplementary Figure 8C). *LGALS1* was found to be expressed by Fibroblasts, Pericytes and Myeloid cells, all of which upregulate its expression in cancer compared to normal tissue (Figure 4D). The hyaluronidase 2 (*HYAL2*) is specifically expressed in Endothelial cells and upregulated in different types of cancer (Figure 4E). Given that most of the GRGs found in Pericytes seem to be present also in Fibroblasts, we focus on the genes specifically found in the latter to better characterize its contribution to the overall expression of GRGs in the tumor. Interestingly, most of the GRGs that characterize Fibroblasts (as *HEXA/B, CHSY1, CSGALNACT1/2, CHPF* and *DSEL*) are associated with proteoglycan metabolism, specifically of chondroitin and dermatan sulfate. To confirm the specific expression of this pathway in Fibroblasts, we calculated a score for the gene set GO term “Chondroitin sulfate - Proteoglycan biosynthetic process” in the scRNA-seq integrated dataset. We found that this pathway is are greatly associated with Fibroblasts, which they upregulate in the tumor compared to normal tissue (Figure 4F). Our data shows that stromal and immune cells can also contribute to the expression profile of GRGs and GRPs present in tissue, but in a rather conserved manner between the various malignancies analyzed.

### Cancer cell lines partially reflect the tumor glycosylation patterns found in tissue

Cancer cell lines are *in vitro* tools that have been widely used for basic research, drug discovery, and glycomic analysis. As cancer cells are the main contributors of the specific tumor glycosylation signatures, we next analyzed if these patters can be recapitulated in cancer cell lines. For this, we used four transcriptomic data sets that collectively represent 1489 cell lines, that were individually clustered using the GRGs (Figure 5A)^30-33^. Given that the same cell line could be clustered different according to datasets, we performed a cluster of clusters as previously described, which allowed us to find consensus clusters while conserving the diversity of the individual data sets (Supplementary Table 4, Supplementary Figure 9A). This strategy led to the identification of five GRG expression clusters in cancer cell lines, which were further characterized by performing gene set enrichment analysis (GSEA) and differential gene expression (Figure 4A-C, Supplementary Figure 9B-E).

Two of the clusters are associated with specific organs and represent the cell lines derived from *Melanoma* and *Hematological* cancers. These clusters are enriched with the gene sets associated with pigmentation and immune responses, respectively (Supplementary Figure 9D). Similar to our findings tissue, melanoma cells lines are enriched in the expression of genes associated with metabolism of sialic acid, including the sialyltransferases *ST3GAL4* and *ST3GAL6* (Figure 5D). The *Neuroendocrine* cluster is associated with neurotransmission gene sets and is mainly composed by cell lines from small cell lung cancer, neuroblastoma and Ewing sarcoma (Figure 5B, Supplementary Table 4). No specific glycosylation pathway was associated with this cluster, although they express the gene *FUT9*, which was overexpressed in Brain cancers in bulk transcriptomic analysis (Figure 5D, Supplementary Figure 2). We also found two heterogeneous clusters that contain cell lines derived from different organs, which we named as *Epithelial* and *Mesenchymal* given that they are enriched in gene sets associated with cell-to-cell contact and epithelial to mesenchymal transition, respectively (Supplementary Figure 8D). Our results show that *Epithelial* cell lines are particularly enriched in the expression of transcripts and gene transcript sets associated with mucin-type *O*-glycosylation and fucosylation (Figure 5D, Supplementary Figure 9E). The most overexpressed gene transcript in this cluster was *GALNT3*, which has been linked to maintaining the epithelial phenotype in different contexts^34,35.^ On the other hand, *Mesenchymal* cells are correlated with the expression of genes involved in proteoglycan metabolism and overexpress the gene encoding for Galectin-1 (*LGALS1*, Supplementary Figure 9E).

**Figure 5.**
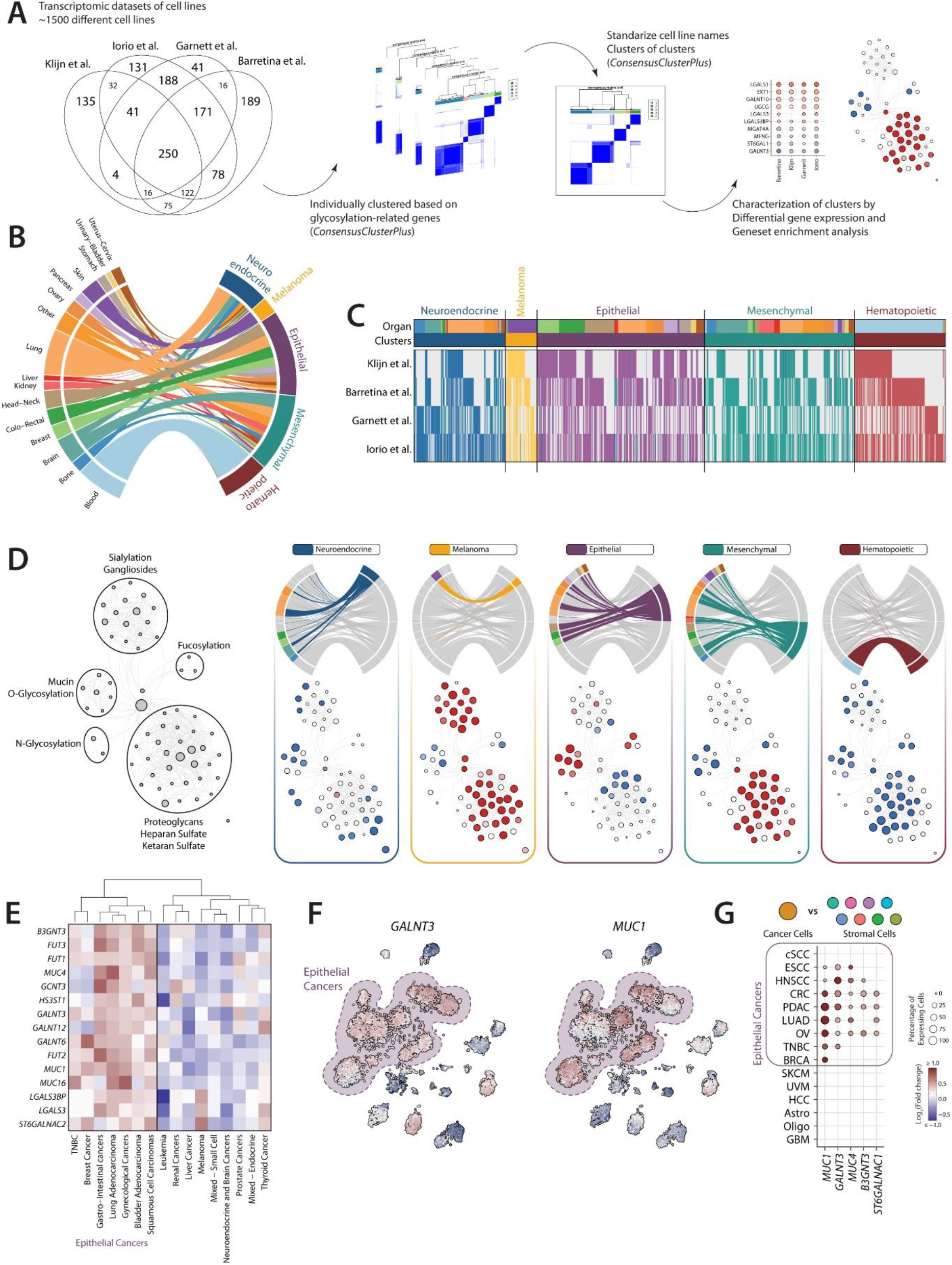
Cancer cell lines partially reflect the GRG expression signatures found in tissue. A) Workflow for the analysis of glycosylation clusters in cancer cell lines. B) Chord diagram showing the association of the glycosylation clusters with the organ of origin of the cell lines. C) Heatmap of cell lines membership to each glycosylation cancer in the different data sets. D) Network representation of the gene set enrichment analysis results using Enrichment Map. E) Heatmap of GRGs associated with *Epithelial* cell lines in the different tissue meta-clusters. F) Expression of *GALNT3* and *MUC1* in tissue samples in the TCGA dataset. G) Dot plot representing the differential expression of GRGs associated with epithelial cell lines in cancer cells vs stromal cells scRNA-Seq. Only significative differentially expressed genes are displayed.

These data suggest that cancer cell lines present a simplified landscape of GRGs compared to the ones found in tissue. However, given that our results can be influenced by the tumor types present in each data set and the difference in the number of cell lines and tissue analyzed, we investigated if similar patterns of GRGs expression can also be found in the TCGA data set. For this, we generated metaclusters by performing hierarchical clustering using the median expression of each GRGs in the clusters identified in Figure 1. Two main metaclusters were found, one of which contains most of the cancers classically associated with an epithelial phenotype (Gastro-Intestinal, Lung, Gynecological, Breast and Squamous cell cancers; Supplementary Figure 8F). These results are in line with our previous analysis, which showed that the cell lines derived from Pancreas, Colorectal, Stomach, Head and Neck, Ovary and Breast cancers are grouped together with the Epithelial cluster, while the ones from Kidney, Liver and Brain cancer presented a mesenchymal phenotype (Supplementary Figure 9C). The Epithelial tissue metacluster showed an expression profile similar to the Epithelial cluster found in cell lines, including *GALNT3* and *MUC1* (Figure 5E-F). Similar results were found in the analysis of single cell transcriptomics (Figure 5G). We next checked if cell lines can still show the expression of tissue-specific GRGs, despite not forming an independent cluster (Supplementary Figure 9G-L). Similarly, to the profiles identified in tissue, we observed the expression of terminal fucosylation genes in cell lines from Gastro-intestinal cancer, *HAS3* and *LGALS7* in SCC and *MUCL1* in non-TNBC Breast cancer (Supplementary Figure 9G-J). However, we did not observe the association of *ST6GAL1* with Liver cancer or *ST8SIA1* with Brain cancers (Supplementary Figure 9K-L). Together, these findings show that cancer cell lines only partially recapitulate the profile of GRG observed in tissue.

### Association of GRG expression patterns with the clinical outcome of patients

We next investigated if the expression of different glycosylation-related genes and pathways was associated with clinical outcome. To analyze this, we performed a systematic survival analysis in the TCGA data set by grouping the patients based on the GRGs and GRPs expression in the top or bottom third and studying the difference in survival rate between these groups using Kaplan-Meier curves and statistical analysis using Cox proportional hazard model (Figure 6A). This analysis was repeated for each combination of glycosylation cluster and TCGA project (Figure 6A-B, Supplementary Figure 10).

**Figure 6.**
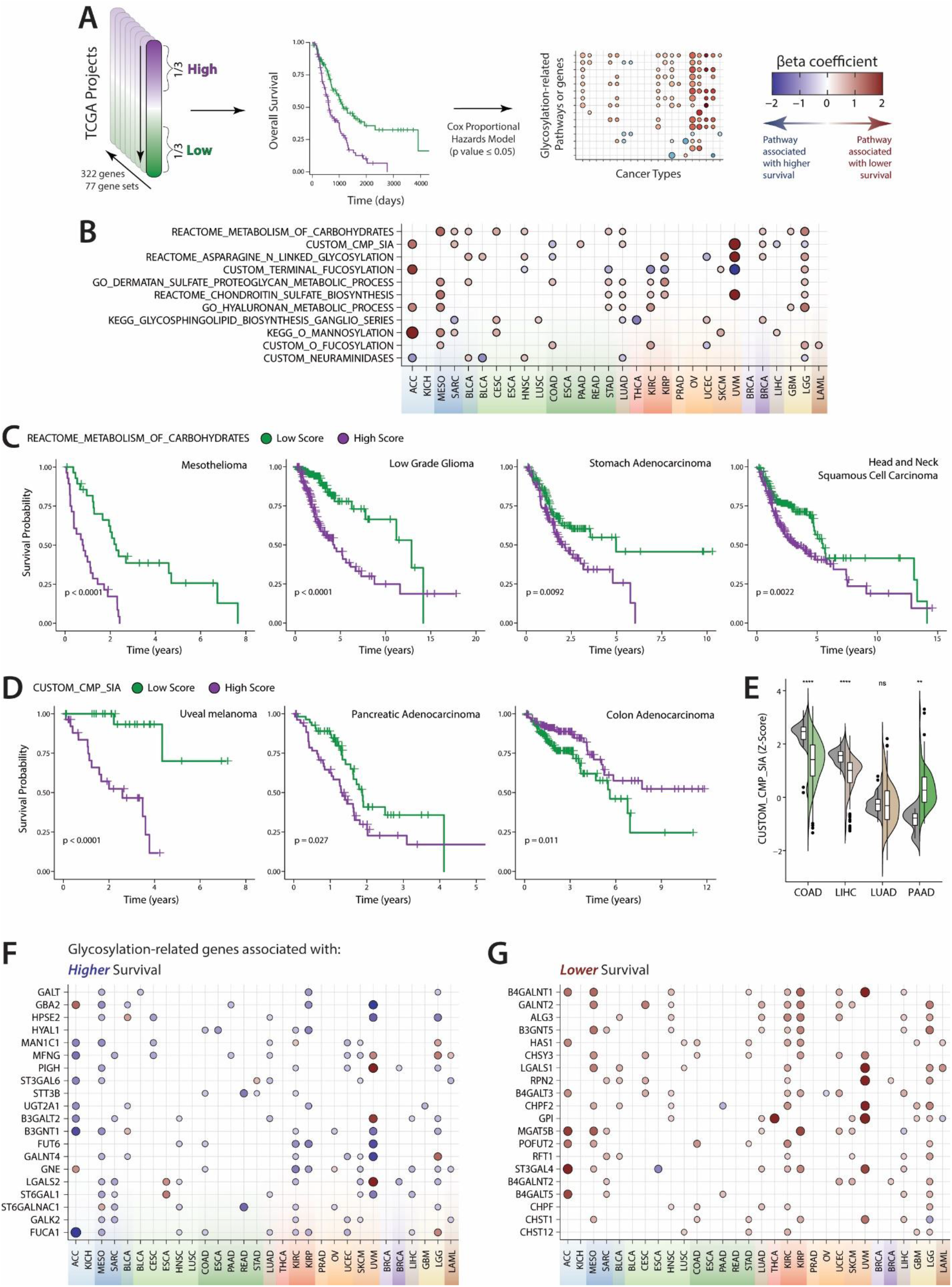
Association of different glycosylation pathways with clinical outcome of patients. A) Workflow used to study the association of glycosylation-related genes and pathways with survival. B) Dot plot summarizing the association of glycosylation-pathways with survival. Kaplan-meier curves illustrating the survival of patients that present a high or low score for the Reactome gene set “Metabolism of Carbohydrates” (C) and the custom gene set “Sialic acid – CMP synthesis” (D) in different tumor types. Statistics: Log-rank test. E) Expression of the custom data set “Sialic acid – CMP synthesis” in normal (gray) and tumor (color) tissue. Statistics: Wilcoxon test (*p ≤ 0.05; **p ≤ 0.01; ***p ≤ 0.001; ****p ≤ 0.0001). In violin plot: data presented as boxplot indicate the median, 25th and 75th percentiles (hinges) and whiskers represent 1.5 times the interquantile range. Dot plot of summarizing the results of the survival analysis of glycosylation-related genes mostly associated with higher (F) or lower (G) survival.

The glycosylation pathway associated with more aggressive cancer was the Reactome gene set “Metabolism of carbohydrates”, in agreement with previously published reports (Figure 6B-C)^36^. This gene set includes a great variety of genes encoding for glycoproteins, transporters, and enzymes related to glycosylation and general carbohydrate metabolism, suggesting that a general upregulation of the pathways associated with glycan metabolism is associated with tumor progression (Figure 6B-C). Pathways corresponding to the biosynthesis of different glycosaminoglycans (dermatan sulfate, chondroitin sulfate and hyaluronan) were also associated with a lower survival of cancer patients (Figure 6B). This may reflect the role of fibroblast activation in cancer progression, given that glycosaminoglycans-related genes are mainly expressed in this cell type and upregulated in tumor tissue (Figure 5F).

The association of some glycosylation pathways with survival depends on the cancer type, as is the case for “*Synthesis of CMP-Sialic acid*”, “*Terminal fucosylation*” or “*Ganglioside biosynthesis*” (Figure 6B). For example, the expression of the gene set reflecting the “*Synthesis of CMP-Sialic acid*” is associated with better survival in Colorectal and Liver cancer, but with lower survival in Sarcoma, Uveal melanoma and Lung, Pancreatic and Breast cancer (Figure 6B-D). Remarkably, this seems to be related to the expression in normal tissue (Figure 6E). This pathway is associated with better survival in cancers of organs where is highly expressed in normal tissue and present a downregulation in cancer samples (as colon and liver), while the opposite is observed in pancreatic cancer (Figure 6E). We repeated this analysis to study the correlation of genes with survival, which led to the identification of a series of genes whose expression is associated with lower or higher survival in different cancer types (Figure 6F-G). Intriguingly, some of the genes associated with a better survival are associated with different immune populations, as *MFGN* and *LGALS2* (Figure 4C), suggesting that these results reflect the anti-tumor immune responses (Figure 6F). On the other hand, some glycosylation-related genes associated with lower survival can be related to the induction of regulatory immune responses, as is the case of LGALS1 (known by its immunomodulatory properties) and ST3GAL4 (one of the enzymes responsible for the synthesis of Siglec-9 ligands)^9,37.^ In conclusion, this analysis allowed us to identify associations of glycosylation-related genes and pathways with survival, showing that some association seem to be conserved between different tumor types, while others are highly dependent with the tumor type.

## Discussion

In this paper we performed a deep characterization of the transcriptional landscape of GRGs and GRPs in a pan-cancer setting. The clustering of samples based on GRGs revealed patters of expression highly dominated by the tissue of origin and tumor type, in a similar way that previous pan-cancer studies^10^. However, we also observed shared patterns of GRGs among different tumor types, as is the case for gastro-intestinal cancers and squamous cell carcinomas, which cluster together independent of the organ of origin. Moreover, our results show that TNBC cluster separated from other types of Breast cancer, illustrating their unique GRGs expression patterns.

The identification of different GRGs and GRPs enriched in different glycosylation clusters, can serve as a source for the discovery of new diagnostic biomarkers. This is supported by the fact that some of the genes and pathways identified in our analysis are associated with biomarkers currently used in the clinic, as is the case of MUC16 for Gynecological cancers or CA19-9 (sialyl Lewis^A^, a fucosylated antigen) for Gastro-Intestinal cancers^3,6,7^. In this paper, we studied as a proof of concept the potential use of serum levels of Galectin-7 for the differential diagnosis of NSCLC, after observing increased levels in Squamous cell carcinoma but not in Adenocarcinoma. Although studies in larger cohorts of patients are needed to confirm our results, our results represent a promising resource for biomarker discovery. The same idea can be used for other tissue specific genes as Galectin-4 (*LGALS4*) for Gastro-Intestinal cancers, *MUC21* for Lung adenocarcinoma, *MUC15* for Thyroid cancer and *MUCL1* for Breast cancer.

Given that several tumor-associated glycans have been associated with the induction of a tolerogenic microenvironment, the identification of specific GRGs and GRPs can aid to better understand glycan-mediated regulatory programs in cancer^2,38.^ Gastro-intestinal cancers are highly enriched in the expression of fucosyltransferases responsible for the synthesis of Lewis antigens, suggesting that these tumor types may be particularly affected by the fucose-specific immunomodulatory receptor DC-SIGN (Dendritic Cell-Specific Intercellular adhesion molecule-3-Grabbing Non-integrin). Triggering of DC-SIGN on antigen presenting cells by fucose-containing glycans induce the expression of IL-10 and enhance the capacity to differentiate Th2 cells^35,39.^ DC-SIGN ligands present in colorectal cancer cell lines suppress dendritic cell function and differentiation^40,41.^ In pancreatic cancer, the expression of fucosylation-related genes by cancer cells is correlated with a tolerogenic phenotype in macrophages^35^. Moreover, the presence of DC-SIGN+ macrophages are associated with a suppressive microenvironment and poor clinical outcome in gastric cancers^42^. Brain cancers are characterized by a high expression of the α1-3 fucosyltransferase FUT9, which has been shown to catalyze the synthesis of the Lewis^X^ antigen in mouse models^43^. We previously reported that human myelin is decorated with Lewis^X^ antigens that mediate the interaction with DC-SIGN+ microglia, inducing a immunoregulatory program that contribute to the immune homeostasis in healthy brain^44^. However, its expression in brain cancer may contribute to the tolerogenic microenvironment.

Sialic acid is another glycan known for its immunomodulatory properties and has been described as a self-associated molecular pattern^45-48^. They are linked to the underlying glycan structures in different linkages (namely α2-3, α2-6 and α2-8), each of which is catalyzed by a group of different enzymes^45^. Here we describe that enzymes that catalyze α2-3 sialylation are associated with Melanoma, α2-6 sialylation with Thyroid cancer and α2-8 sialylation (also called polysialylation) within brain cancers. Sialylated glycans can be recognized by a family of lectin receptors called SIGLECs (sialic acid-binding immunoglobulin-like lectins), most of which present a ITIM (immunoreceptor tyrosine-based inhibitory motif) sequence in their intracellular domain^2,45.^ It was previously reported that ligands of Siglec-7 and Siglec-9 can be found in cancer cells and cell lines from different tumor types, but are particularly increased in Melanoma^46^. This is in line with our results that Melanoma tissue samples and cell lines highly express *ST3GAL4*, which is shown to synthesize the ligands of Siglec-9^37,47.^

The use of scRNA-Seq data sets allowed us to resolve the expression of GRGs at a single cell level, identifying cancer cells as the main contributors to the specific genes observed in the analysis of bulk transcriptomic data. Our results suggest that the expression dynamic of GRGs and GRPs in tumor development are influenced by the cell of origin and the oncogenic pathways that lead up to malignant transformation. For example, pancreatic and colorectal cancers cells, despite having a similar expression profile of GRGs and GRPs, present different changes with respect to their normal counterparts. We hypothesize that KRAS mutations can push cells to a similar glyco-phenotype independent of the cell of origin, given that it is an oncogenic driver commonly found in both cancer types^28^. It has been previously shown that KRAS mutations can lead to different signaling pathways and transcriptomic changes in a tissue-specific manner^49,50.^ The complexity of oncogenic pathways that are present in cancer hinders the systematic study in a pan-cancer setting and highlights the need to study their effect in the expression of GRGs and GRPs in the context of specific organs or cancer types, considering also clinically relevant variables.

Despite that the cancer cells seems to be responsible for the expression of specific GRGs and GRPs in each tumor type, stromal cells can also contribute to the overall tumor glyco-code, although in a rather conserved form across different tumor types. For example, Galectin-1 (*LGALS1*) was mainly expressed by Fibroblasts, Pericytes and Myeloid cells, all of which upregulate its expression in tumor compared to normal tissue. Stromal-derived Galectin-1 has been associated with vascularization and metastasis formation in pancreatic cancer, proving to be a good candidate for therapeutic targeting^51^. Tumor-associated endothelial cells appear to increase the expression of Hyaluronidase 2 (*HYAL2*), one of the enzymes responsible for the degradation of hyaluronan, the main component of the endothelial glycocalyx^52^. This process leads to the generation of the pro-tumoral mediator low-molecular-weight hyaluronan, which can induce inflammation, immune cell recruitment and promote migration of cancer cells^53,54.^ The characterization of the stromal contribution to the overall profile of GRGs and GRPs found in tumor tissue is essential for the correct interpretation of the results obtained with the analysis of bulk transcriptomics.

We also showed that cell lines present simplified glycosylation patterns compared to the respective tumor tissue. Cell lines are an important tool in cancer research, although several studies have shown that they differ in their ability to reflect the primary tumor^55-58^. The clustering of cell lines based on GRGs led to the identification of two general clusters, Epithelial and Mesenchymal, that correspond to most of the cell lines and three tissue-specific clusters (Melanoma, Neuroendocrine and Hematopoietic). Of these clusters, Melanoma cell lines seems to represent better the GRGs and GRPs present in tissue, characterized by a high expression of sialylation-related genes. Klijn *et al* have shown that the unsupervised clustering of cell lines is dominated by the Epithelial-to-mesenchymal transition (EMT), which is in line with our results^32^. Epithelial cancer cell lines and tissues are characterized by the expression of genes associated with Mucin-type O-glycosylation and terminal Fucosylation, similarly to our prior results in pancreatic cancer^35^. One of the genes associated with these cell lines is *GALNT3*, which has been associated with the epithelial phenotype in different contexts^34,35.^ Conversely, the fucosylation of TGFβ receptors in colorectal cancer cell lines by *FUT3* and *FUT6* is essential for the TGFβ-mediated induction of EMT, which suggest that enrichment of fucose-related genes in epithelial cells “prepare” these cells to undergo to this transition^59^. In Mesenchymal cells we found an enrichment of Galectin-1 (*LGALS1*), which has been associated to EMT in hepatocellular carcinoma^60^. Therefore, there is a need for a rigorous selection of the cell lines to be used in the study of GRGs and GRPs in cancer biology, based on their capacity to reflect the expression of the gene of interest in tissue. However, it is also important to study the if cancer cells in tissue present different glycosylation subtypes that can be reflected in different cell lines. We previously reported in that pancreatic cancer cell lines partially reflect the different glycosylation profiles found on the cancer cells present in the Basal and Classical subtypes^35^.

Finally, we identified GRGs and GRPs associated with the clinical outcome of patients, which indicate that changes in the tumor glyco-code affects disease progression. The transcriptomic analysis of GRGs had led to the identification of tumor subtypes with differences in the overall survival in pancreatic and breast cancer^35,61.^ Interestingly, our analysis showed that the association of several pathways with survival depends on the tumor type, as is the case for “*Synthesis of CMP-Sialic acid*”, which leads to the generation of the glycan donor used by sialyltransferases. This pathway is associated with a worst survival uveal melanoma, pancreatic and lung cancer, while the opposite is observed in colorectal and liver cancer. We previously reported that the knockout of CMAS, a key enzyme in this pathway, leads to a more aggressive tumor in a mice model of colorectal cancer^62^.

The characterization of the differential contribution of the tumor and stromal compartment to the overall expression profile allows a more accurate interpretation of the results obtained from the analysis of bulk transcriptomics. Several of the GRGs and GRPs commonly correlated with survival in different tumor types are actually associated with specific cells in the stromal compartment (as *MFNG* in all immune cells, *LGALS2* in myeloid cells, *ST6GAL1* in B and plasma cells and *CHPF* and *ST3GAL4* for fibroblasts). This may reflect the presence of different cell populations when analyzing survival in bulk transcriptomics and does not necessarily reflect the role of the specific gene or pathway. The use of single cell and spatial techniques for the characterization of tissue samples can help to circumvent this issue and to full characterize the tumor glyco-code. However, the presence of different clinical parameters that can influence the clinical outcome of patients emphasizes the need to perform a characterization of the tumor glyco-code in individual cancer types.

The transcriptional analysis of GRGs and GRPs is a powerful tool for a better understanding of glycosylation and different approaches have been taken to obtain novel insights of its role in health and disease^4,35,63^. However, its potential to predict glycan structures presents several limitations associated with the organization, complexity and regulation of the glycosylation machinery. Alterations in tumor glycosylation can be the result of multiple mechanisms other than the expression of GRGs, including changes in localization of glycosyltransferases or differences in chaperone function^64^. The Tn antigen, one of the most studies glycan structures in cancer, corresponds to the first step of the mucin-type *O*-glycosylation: a simple *N*-Acetyl-galactosamine bound to a serine, threonine o tyrosine in the peptide backbone of a protein^2,38,65,66^ Its synthesis is catalyzed by a family of twenty enzymes called polypeptide *N*-Acetylgalactosaminyltransferase (encoded by the genes *GALNT*) that differ in their tissue distribution and specificity^67,68.^ However, their presence in tumor cells is difficult to predict based only on transcriptomic analysis as it can be influenced by multiple factors: activity or defects of the extending enzymes (C1GALT1 and ST6GALNAC1) or the COSMC chaperone (C1GALT1C1), overexpression of carriers (like mucins) and GALNT re-localization to the endoplasmic reticulum^69-71^. Moreover, the organization of the glycosylation machinery as an “assembly line” is also difficult the prediction of the glycan structures based on the transcriptomics analysis, as there is need to not only know the expression of the gene of interest, but also all the others that contribute to the synthesis of their potential substrates^72^. This is illustrated in colorectal cancer, where fucosylated glycans have been reported to be present in both normal and tumor tissue, in line with our results that show no changes in the expression of fucosyltransferases^73^. However, differences in the specific fucose-containing structures have been recently reported, with cancer cells characterized by the synthesis of sialyl-Lewis core 2 *O*-glycans^73^. This illustrates the need for a multidisciplinary approach to completely understand the role of glycosylation in cancer, integrating transcriptomic and glycomic analysis with functional and clinical studies. Moreover, future efforts to correlate the presence of glycosylation structures within the whole transcriptome will shed a light on novel regulatory pathways and the potential use of gene expression for the prediction of the tumor glyco-code.

## Materials and Methods

### Data availability

Datasets used in this paper are detailed in Supplementary Table 3. TCGA pan cancer dataset was downloaded from https://gdc.cancer.gov/about-data/publications/pancanatlas. Single cell RNA-Seq datasets were downloaded from the NCBI GEO database (https://www.ncbi.nlm.nih.gov/geo/) with the following accession numbers: GSE160269 (Esophageal Squamous cell carcinoma, ESCC)^25^, GSE144236 (cutaneous Squamous cell carcinoma, cSCC)^19^, GSE132465 (Colorectal Cancer, COREAD)^20^, GSE161529 (Brest Cancer, BRCA)^26^, GSE120575 (Melanoma, SKCM)^15^, GSE115978 (Melanoma, SKCM)^16^, GSE139829 (Uveal melanoma, UVM)^22^, GSE146026 (Ovarian cancer, OV)^21^, GSE131928 (Glioblastoma, GBM)^17^, GSE131907 (Lung adenocarcinoma, LUAD)^23^, GSE149614 (Hepatocellular carcinoma, HCC)^24^, GSE103322 (Head and neck Squamous cell carcinoma, HNSCC)^13^, GSE70630 (Low grade glioma – Oligodendroglioma, LGG)^12^ and GSE89567 (Low grade glioma – Astrocytoma, LGG)^14^. For Pancreatic ductal adenocarcinoma (PAAD), Peng *et al*. was downloaded from the Genome Sequence Archive (https://bigd.big.ac.cn/gsa/) under the project PRJCA001063^18^. The TCGA survival data was obtained from the publication of Liu *et al^74^* and the mutational data from Sanchez-Vega *et al^28^.* The datasets with transcriptomic data of cell lines were downloaded from the NCBI GEO database, accession number GSE36133^30^, or from ArrayExpress (https://www.ebi.ac.uk/arrayexpress/), identifiers: E-MTAB-783^31^, E-MTAB-2706^32^ and E-MTAB-3610^33^. The remaining data are available within the Article, Supplementary Data file or available from the authors upon request.

### Analysis TCGA dataset

In the pan-cancer TCGA dataset, we filtered out the low expressed genes by discarding the ones that present less than 5 counts in half of the samples. To analyze the presence of glycosylation profiles hared between the different cancer types, we used package *ConsensusClusterPlus*^*75*^ to perform a clustering using the median-centered expression of the 100 most variable glycosylation genes (defined by interquantile range, IQR) and the following settings: 500 iterations with 90% of sampling, “*hclust*” as method and “*ward*.*D2*” as inner and final linkage. For the gene set enrichment analysis, we used the method *ssGSEA* of the *GSVA* package^76^. We used Receiver Operating Characteristic (ROC) curves to define genes and pathways associated with each cluster, calculating the prediction power from the area under the curve (AUC) as absolute(AUC-0.5)*2. Genes and pathways with a prediction power equal or higher of 0.85 were selected as specific for each cluster.

For the systematic analysis of the association of gene expression with overall survival of patients, we selected the combination of TCGA Project with glycosylation clusters that present at least 60 patients. To analyze the association of different genes and pathways with overall survival, samples were divided in thirds based on gene expression and differences in the survival between top and bottom third was analyzed using the function *coxph* from the package *survival*, using the Cox proportional hazards regression model. Kaplan–Meier curves were generated using the function *ggsurvplot* (from the R package *survminer*).

### scRNA-Seq analysis

*Seurat(v4)* package was used for the analysis of single cell RNA-Seq^27^. For each dataset, we generated Seurat objects with samples obtained from tumor tissue, downsampled to 10.000 cells and proceed to their integration using Fast integration using reciprocal PCA (RPCA) method, following the recommendations of the developers. Clustering was performed using the functions *FindNeighbors* using the first 30 Principal Component Analysis (PCA) dimensions and *FindClusters* with resolution set as 1. Clusters were visualized using uniform manifold approximation and projection (UMAP), using the function *RunUMAP*. Cell identification was done based on gene expression and the cell types present in the original publications. In our analysis, we identified a cluster composed mainly of malignant cells and tissue-specific cell types, which we cleaned up to only select cancer cells. For each tumor type, we selected cells corresponding to this cluster and proceed to normalize and perform PCA. Presence of different cell types was analyzed by running the functions *FindNeighbors, FindClusters* and *RunUMAP* using the first 50 PCA components. Cells were discarded based on the expression of different lineage markers, as *PTPRC* for immune cells, *CD3G* and *CD8A* for T Cells, *MS4A1* and *CD19* for B Cells, *AIF1* and *CD14* for Myeloid cells, *LUM* for Fibroblasts, *RGS5* for Pericytes, *CDH5* for Endothelial cells, *PMEL* for Melanocytes, *MOG* and *MBP* for Oligodendrocytes or *GFAP* for Astrocytes. Differential gene expression in the integrated dataset was performed using custom-made function *FindMarkersPerTumorType*, modified from the function *FindConservedMarkers* of the *Seruat* package.

To analyze the differential gene expression between normal and cancer cells, we performed the scRNA-Seq analysis of the datasets from ESCC^25^, cSCC^19^, COREAD^20^, BRCA^26^, LUAD^23^, HCC^24^ and PDAC^18^. Data was normalized and scaled using the *SCTransform* function (regressing out the mitochondrial percentage), and Principal Component Analysis (PCA) was performed with the 3000 most variable genes. PCA components 1 to 30 were used for graphical based clustering at a resolution of 1. These clusters were projected onto Uniform Manifold Approximation and Projection (UMAP) dimensional reduction. Differential gene expression between normal and cancer cells was done using the function *FindMarkers*. For this analysis, we used epithelial cells in normal and tumor tissue for ESCC, COREAD, PDAC, LUAD and BRCA (identified by the expression of epithelial and cytokeratin genes, as *KRT19, EPCAM* and *CDH1*); Keratinocytes for cSCC (*COL17A1, KRT14* and *KRT10*) and Hepatocytes for HCC (*ALB, FGA* and *FGG*).

### Cell line transcriptomic analysis

Given that the datasets used in this paper sometimes provide slightly different names for the same cell lines, we first uniformized the designation by changing the characters to upper case and eliminating punctuation marks. Moreover, we checked for synonyms of different cell lines using the database Cellosaurus (https://web.expasy.org/cellosaurus/)^77^. This method led to 1489 unique cell lines (Supplementary Table 4).

Each dataset was independently clustered applying the package *ConsensusClusterPlus*^*75*^, using 1000 iterations with 90% of sampling, “hclust” as method and “ward.D2” as inner and final linkage. As an input, the median-centered expression of the 100 most variable glycosylation genes was used. To find consensus clusters between the different datasets, we used cluster of clusters using *ConsensusClusterPlus* as reported before^78^. For characterization of the clusters, we used the dataset from Klijn *et al* to perform Gene Set Enrichment Analysis (GSEA)^79^ as implemented in the javaGSEA Desktop Application and visualized in Cytoscape using the EnrichmentMap app^80^. The *limma*^*81*^ package was used for the differential gene expression analysis between the different clusters.

### Galectin-7 ELISA in plasma samples

Plasma from patients with Lung Adenocarcinoma (n = 20) and Squamous Cell Carcinoma (n = 16) was obtained from the Liquid Biopsy Center of the Amsterdam UMC. The studies involving human participants were reviewed and approved by the Medical Ethical Committee, Amsterdam UMC. Written consent was obtained for all the donors. A Galectin-7 ELISA kit (R&D Systems) was used to measure its concentration in the plasma of patients.

## Code accessibility

The main scripts used in this manuscript can be found in https://github.com/MolecularCellBiologyImmunology/Glyco_PanCancer. Other codes for the transcriptomic analysis are available from the authors upon request.

## Acknowledgements

We would like to acknowledge the work of the research groups responsible for the generation and processing of the datasets used in this paper. We thank Idris Bahce (Department of Pulmonary Diseases, Amsterdam UMC) and the Liquid Biopsy Center (Amsterdam UMC) for their help in obtaining the plasma samples of Lung cancer patients. E.R. is supported by Immunoshape (MSCA-ITN-2014-ETN No 642870) and Spinoza grant (NWO, Nederlandse Organisatie voor Wetenschappelijk Onderzoek). D.L. is supported by KWF 12789-2019. Y.v.K. are supported by the European Research Council (ERC-339977-Glycotreat).

