## Supplementary Data for "The transcriptional landscape of glycosylation-related genes in cancer"

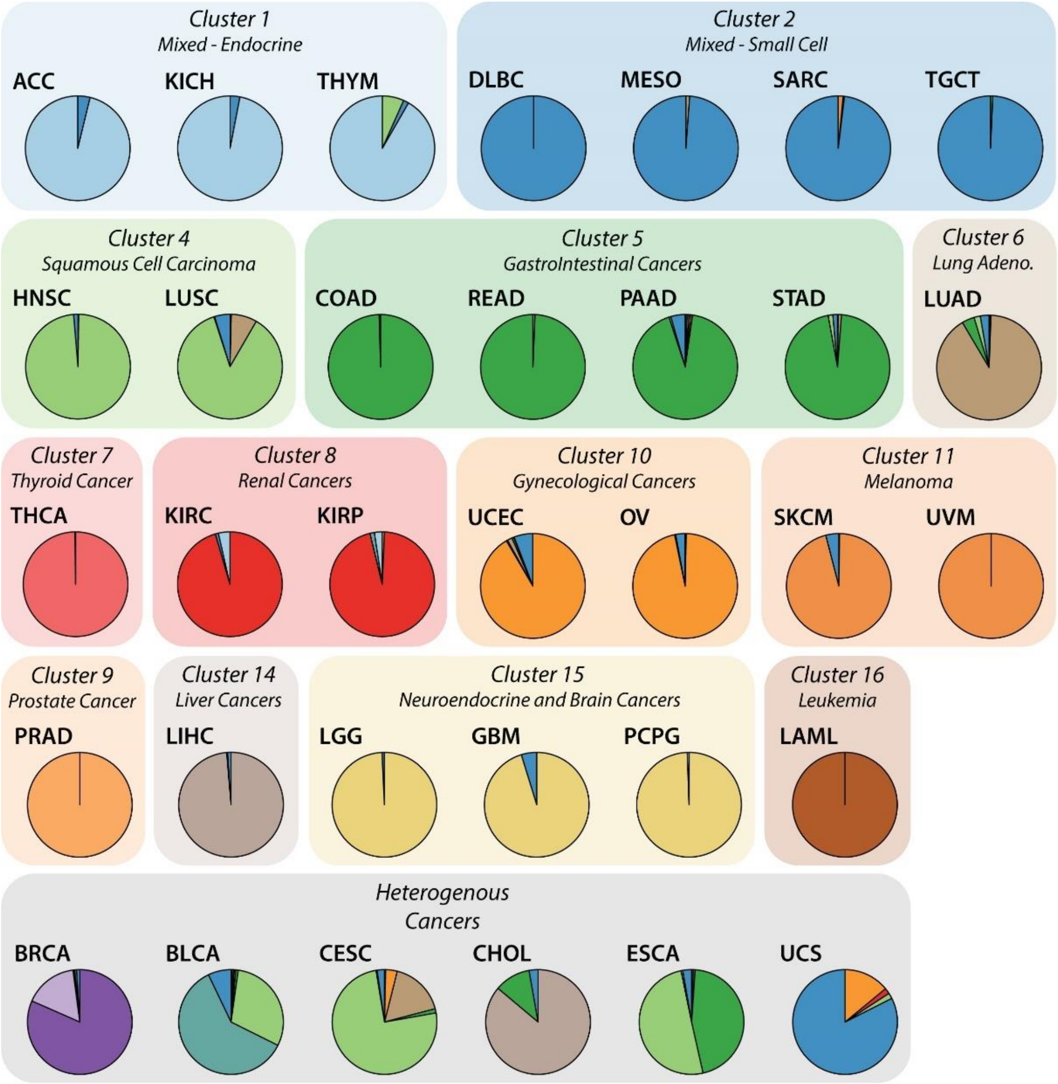


**Supplementary Figure 1. Quantification of the glycosylation clusters associated to a given TCGA project.**

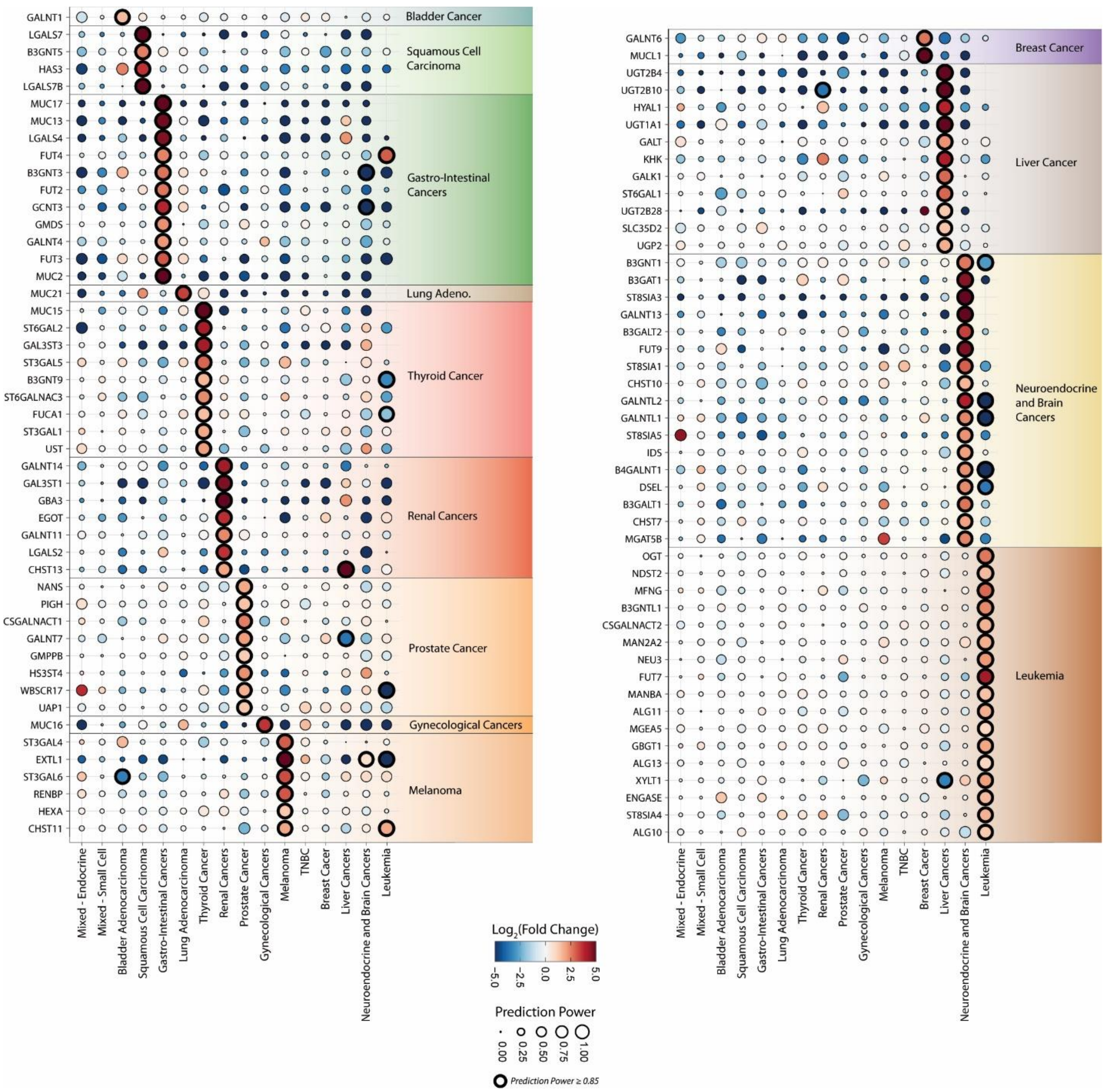


**Supplementary Figure 2. Association of glycosylation-related genes with the different clusters in the TCGA data set.**

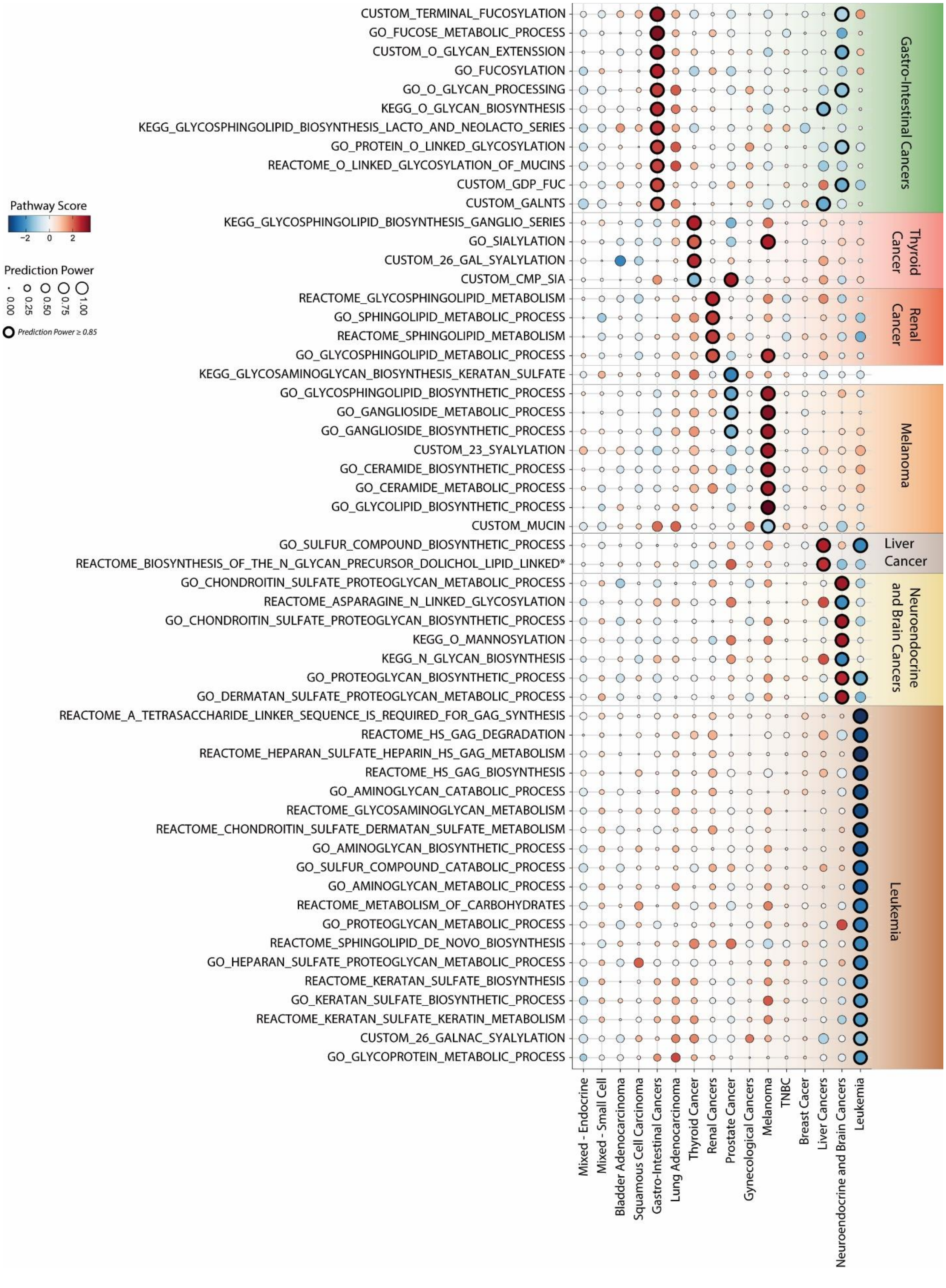


**Supplementary Figure 3. Association of glycosylation pathways with the different clusters in the TCGA data set.** *Full name of gene set: *REACTOME_BIOSYNTHESIS_OF_THE_N_GLYCAN_PRECURSOR_DOLICHOL_LIPID_LINKED_OLIGOSACCHARIDE_LLO_AND_TRANSFER_TO_A_NASCENT_PROTEIN*.

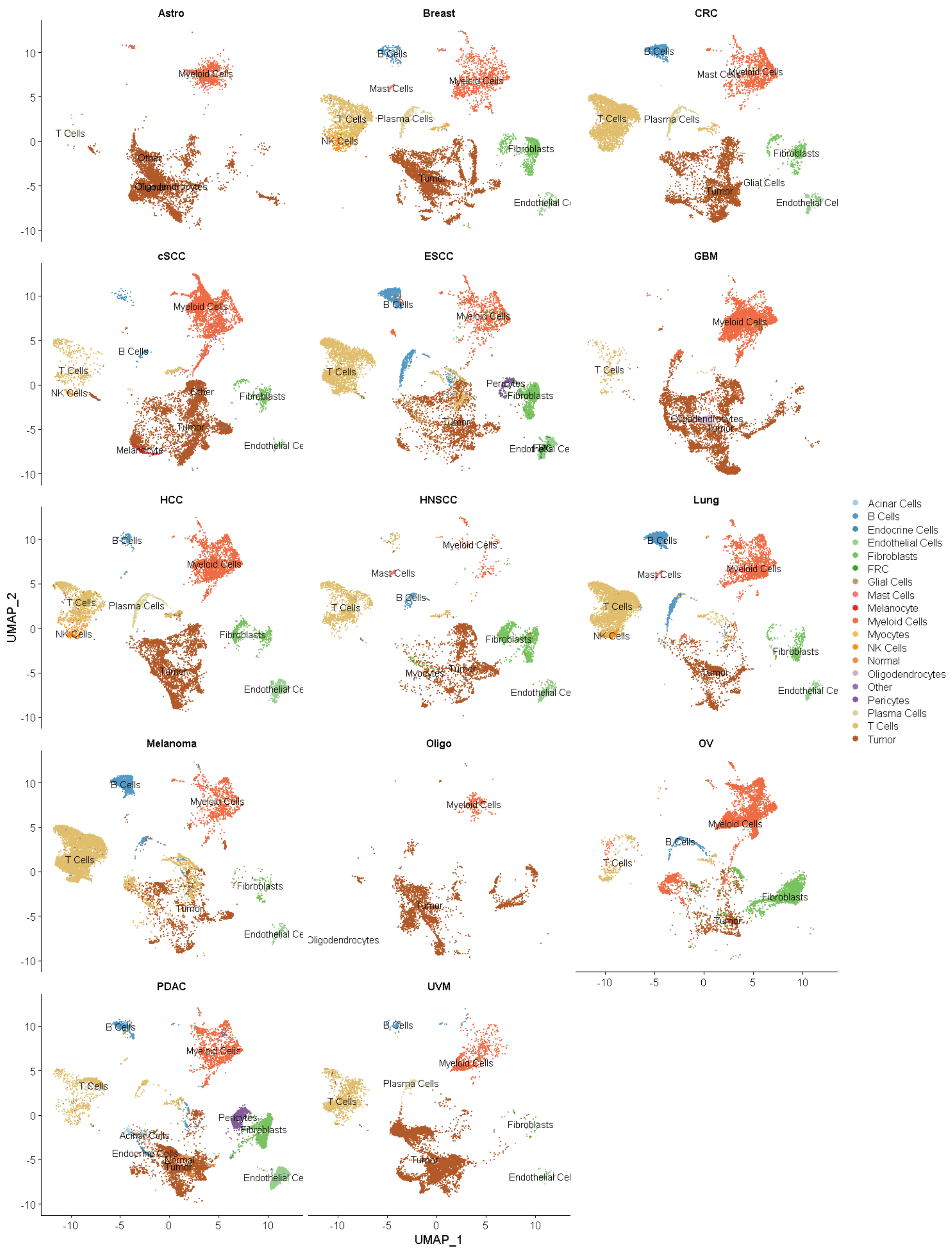


**Supplementary Figure 4. UMAP plot of the integrated single cell RNA-seq dataset displaying the clusters obtained in the independent analysis of each dataset.**

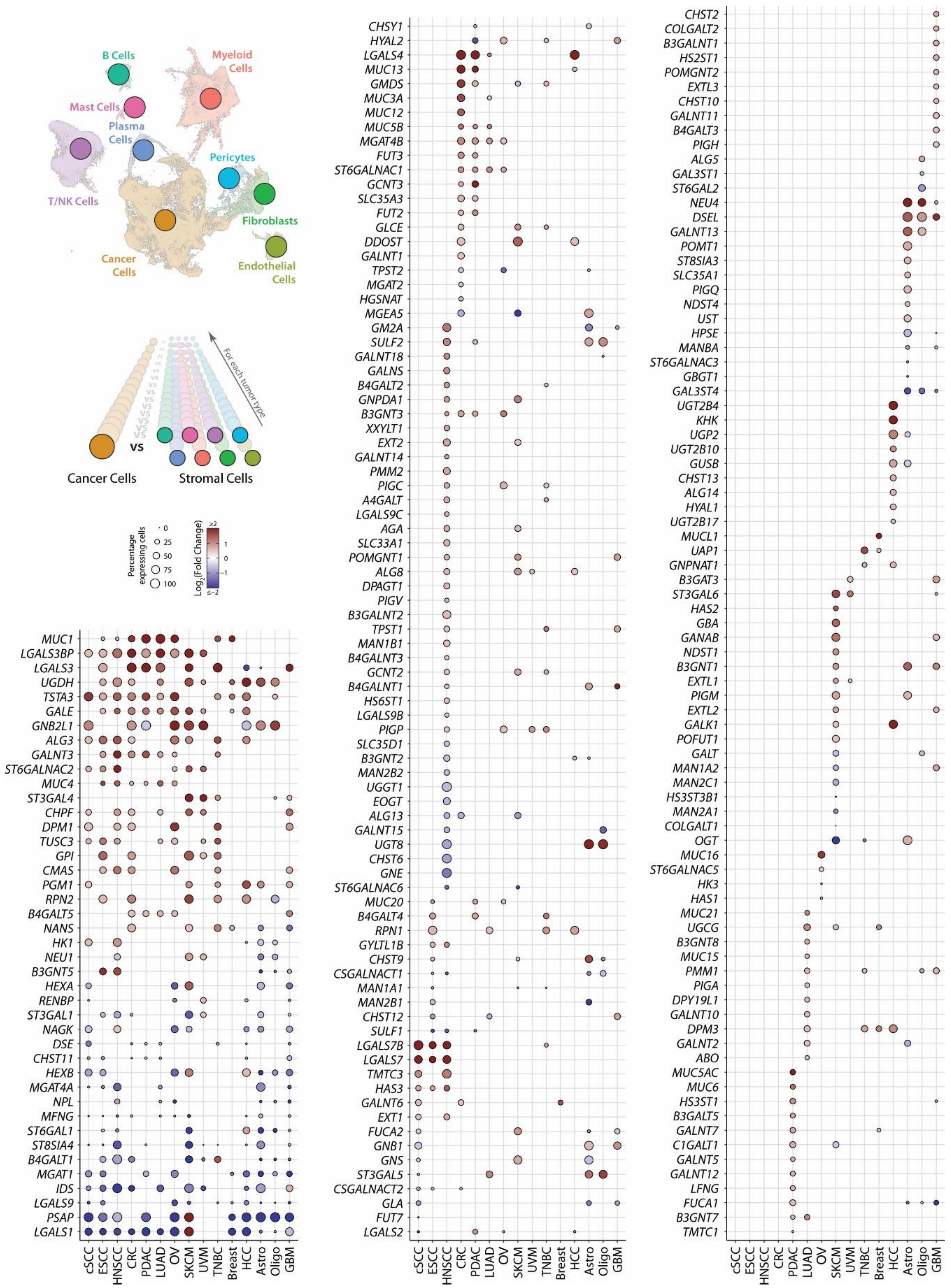


**Supplementary Figure 5. Analysis of the contribution of the cancer and stromal compartment to the expression of GRGs in tumor tissue.** In the integrated data set, we performed differential expression of GRGs in each cell type against all the others in every tumor type individually, using a modified version of the function *FindConservedMarkers* of the package *Seruat*.

**Supplementary Figure 6. Differential expression of glycosylation-related between normal and tumor cells in scRNA-Seq.** A) Top: Expression of specific GRGs in normal and tumor cells in different tumor types. Bottom: Immunohistochemistry staining of different glycosylation-related proteins in normal tissue. Images obtained from the Human Protein Atlas, available from v21.proteinatlas.org, using the corresponding gene names displayed. B) For each dataset, we used the function *FindMarkers* of the *Seurat* package to analyze the differential expression of glycosylation-related genes. The following cell types were used for the analysis between normal and tumor tissue: epithelial cells in in Pancreatic, Colorectal, Lung, Esophagus and Breast cancer, keratinocytes in cutaneous SCC and Hepatocytes in Liver cancer. In violin plot: data presented as boxplot indicate the median, 25th and 75th percentiles (hinges) and whiskers represent 1.5 times the interquantile range.

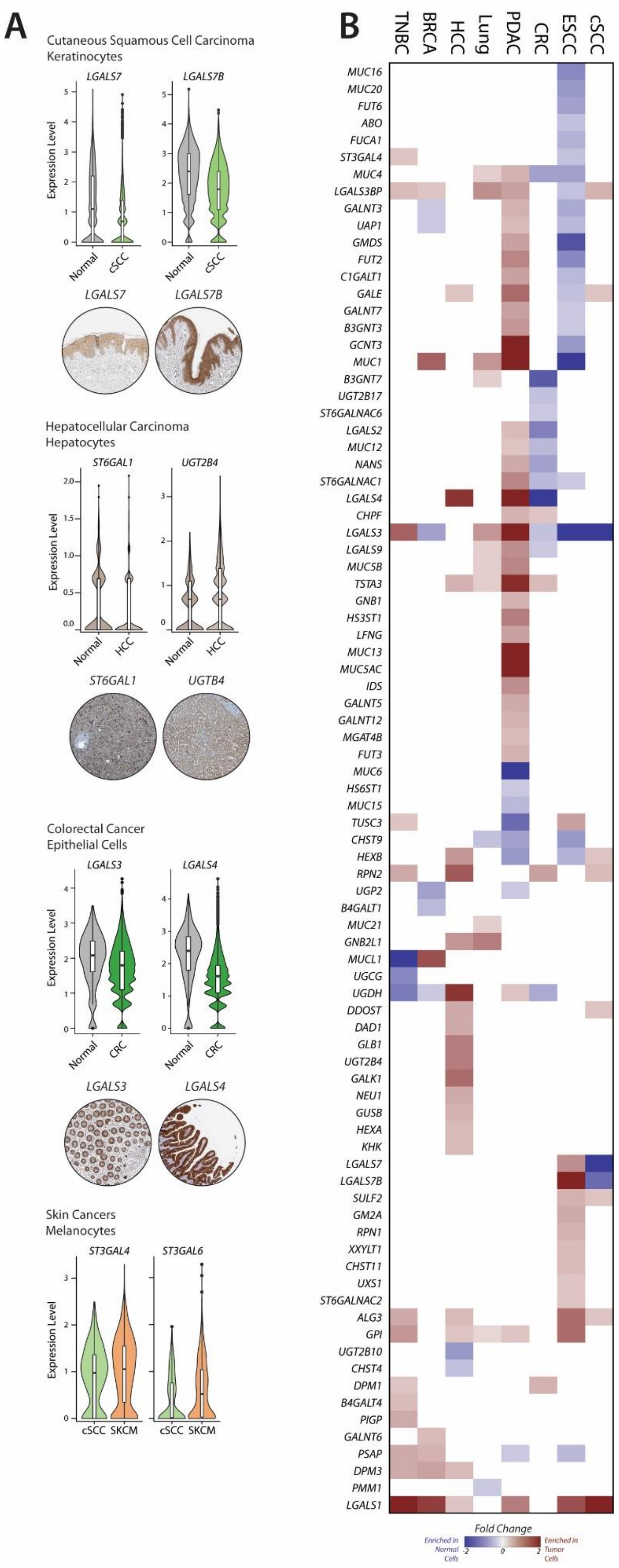

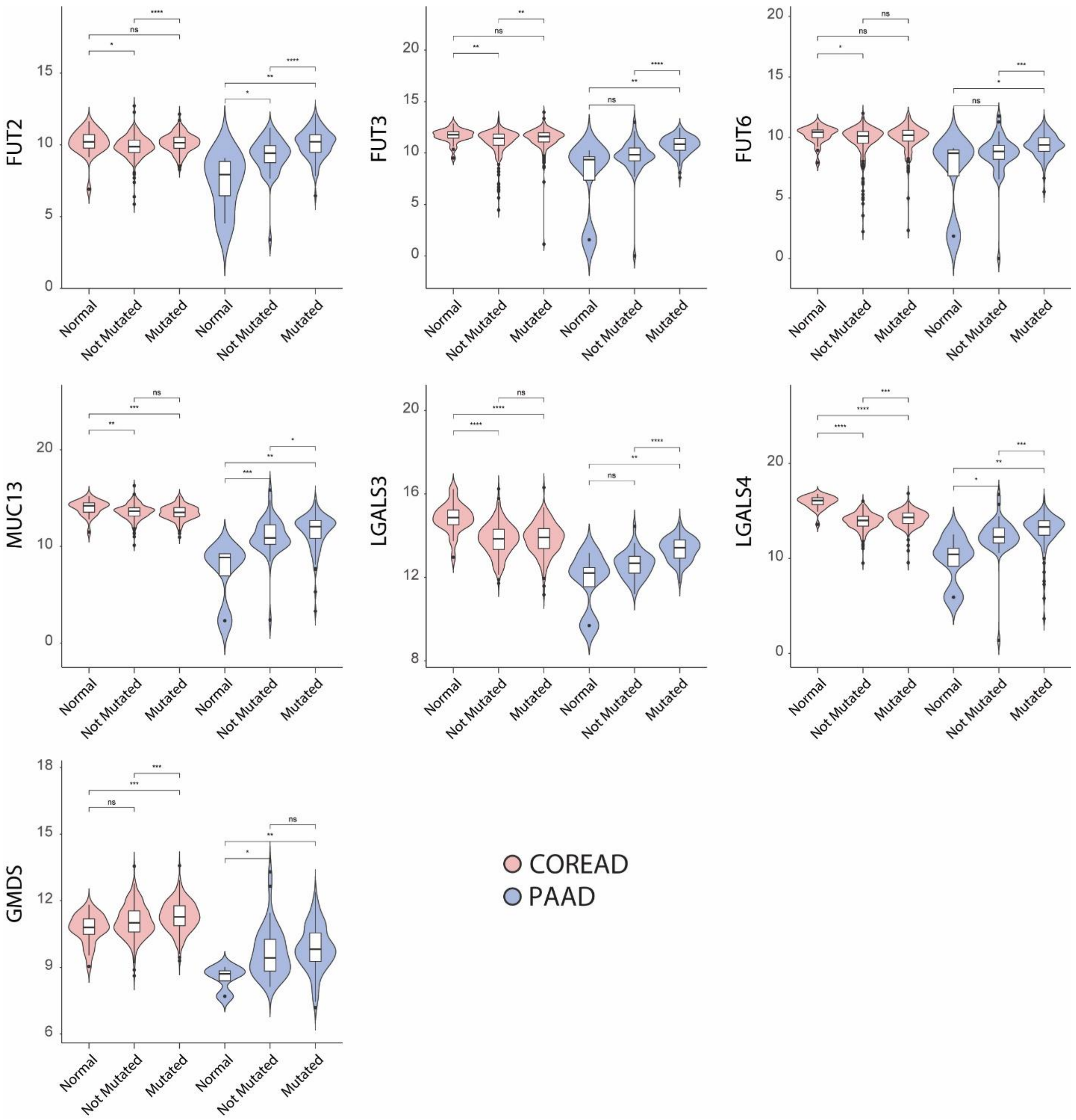


**Supplementary Figure 7. Expression of GRGs associated with gastrointestinal cancers in normal and tumor tissue with or without KRAS mutations.** Samples corresponding to the Gastro-Intestinal cluster were used. In COREAD: Normal (n = 26); Not Mutated (n = 256); Mutated (n = 200). In PAAD: Normal (n=4); Not Mutated (n = 43); mutated (n = 109). Statistics: Wilcoxon test (*p ≤ 0.05; **p ≤ 0.01; ***p ≤ 0.001; ****p ≤ 0.0001). In violin plot: data presented as boxplot indicate the median, 25th and 75th percentiles (hinges) and whiskers represent 1.5 times the interquantile range.

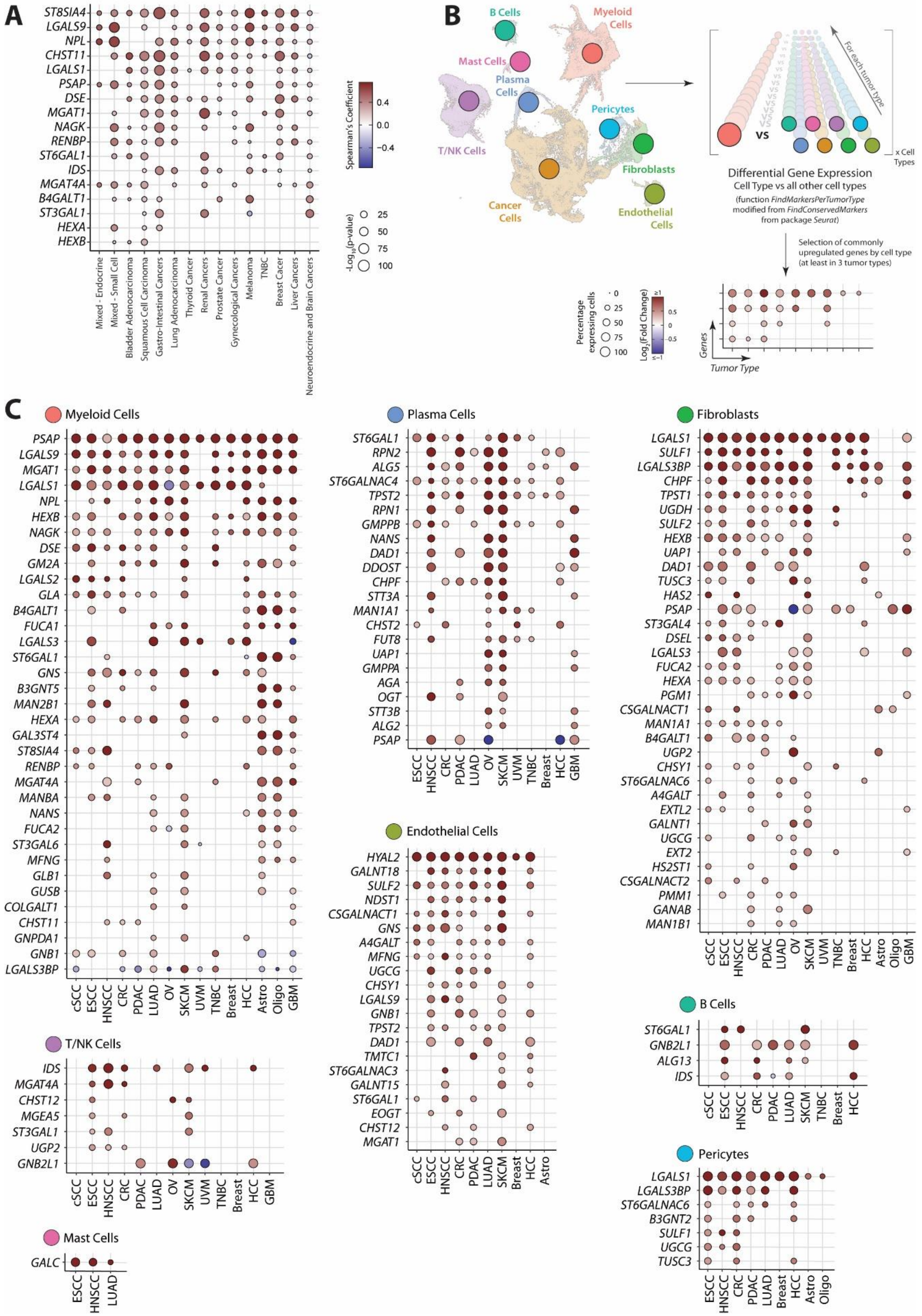


**Supplementary Figure 8. Analysis of the contribution of stromal cells to the expression of GRGs in tumor tissue.** A) Correlation between the stromal fraction in the TCGA dataset and the expression stromal-associated GRGs (identified in scRNA-Seq data, Supplementary Figure 5). B) In the integrated data set, we performed differential expression of GRGs in each cell type against all the others in every tumor type individually, using a modified version of the function *FindConservedMarkers* of the package *Seruat*. C) Dot plot showing the expression of cell specific GRGs.

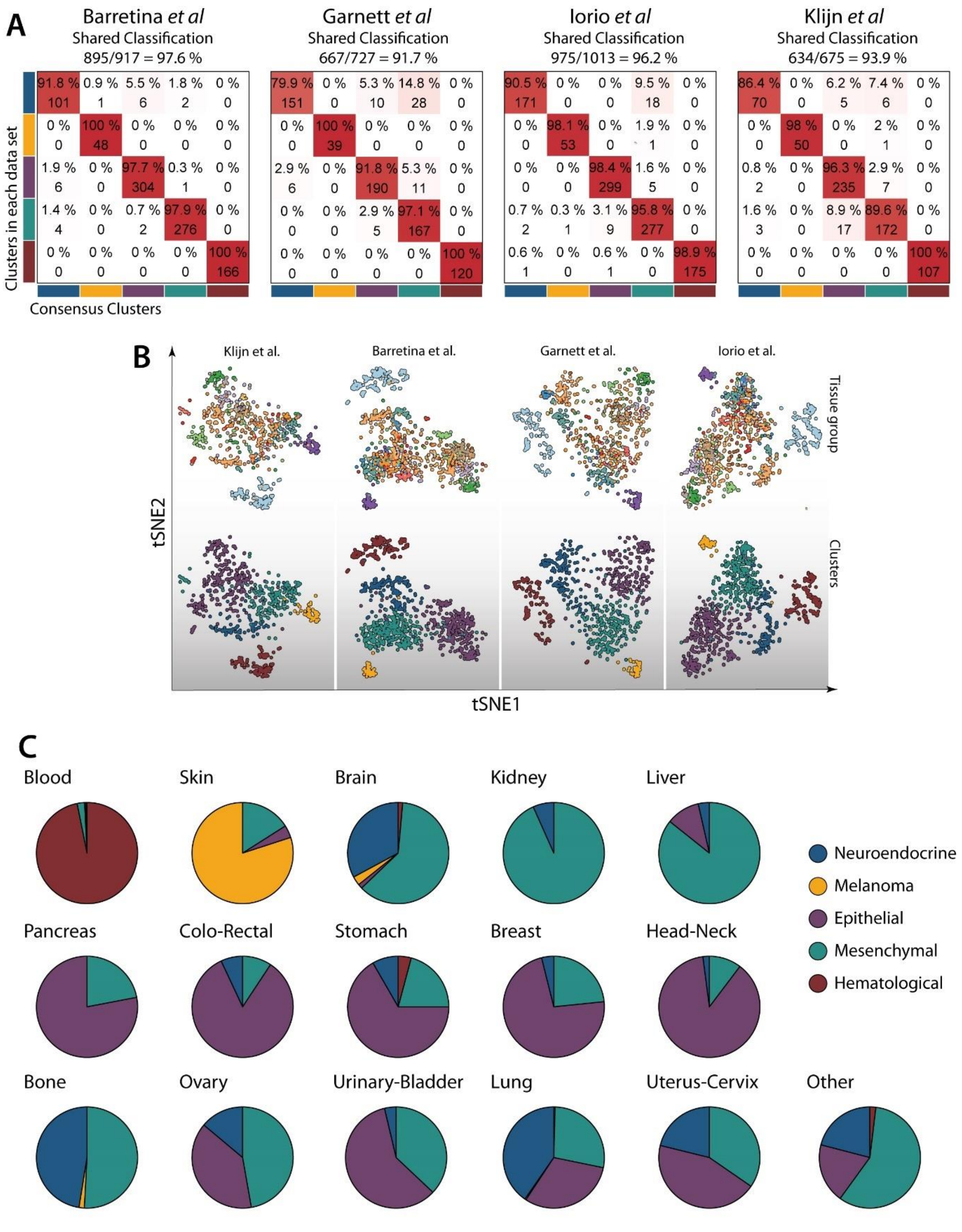
A
Barretina et al
Shared Classification
895/917 = 97.6 %
91.8 % 0.9 % 5.5 % 1.8 % 0 %
101 1 6 2 0
0 % 100 % 0 % 0 % 0 %
0 48 0 0 0
1.9 % 0 % 97.7 % 0.3 % 0 %
6 0 304 1 0
1.4 % 0 % 0.7 % 97.9 % 0 %
4 0 2 276 0
0 % 0 % 0 % 0 % 100 %
0 0 0 0 166
Garnett et al
Shared Classification
667/727 = 91.7 %
79.9 % 0 % 5.3 % 14.8 % 0 %
151 0 10 28 0
0 % 100 % 0 % 0 % 0 %
0 39 0 0 0
2.9 % 0 % 91.8 % 5.3 % 0 %
6 0 190 11 0
0 % 0 % 2.9 % 97.1 % 0 %
0 0 5 167 0
0 % 0 % 0 % 0 % 100 %
0 0 0 0 120
Iorio et al
Shared Classification
975/1013 = 96.2 %
90.5 % 0 % 0 % 9.5 % 0 %
171 0 0 18 0
0 % 98.1 % 0 % 1.9 % 0 %
0 53 0 1 0
0 % 0 % 98.4 % 1.6 % 0 %
0 0 299 5 0
0.7 % 0.3 % 3.1 % 95.8 % 0 %
2 1 9 277 0
0.6 % 0 % 0.6 % 0 % 98.9 %
1 0 1 0 175
Klijn et al
Shared Classification
634/675 = 93.9 %
86.4 % 0 % 6.2 % 7.4 % 0 %
70 0 5 6 0
0 % 98 % 0 % 2 % 0 %
0 50 0 1 0
0.8 % 0 % 96.3 % 2.9 % 0 %
2 0 235 7 0
1.6 % 0 % 8.9 % 89.6 % 0 %
3 0 17 172 0
0 % 0 % 0 % 0 % 100 %
0 0 0 0 107
Clusters in each data set
Consensus Clusters
B
Klijn et al.
Barretina et al.
Garnett et al.
Iorio et al.
Tissue group
Clusters
tSNE2
tSNE1
C
Blood
Skin
Brain
Kidney
Liver
Pancreas
Colo-Rectal
Stomach
Breast
Head-Neck
Bone
Ovary
Urinary-Bladder
Lung
Uterus-Cervix
Other
Neuroendocrine
Melanoma
Epithelial
Mesenchymal
Hematological

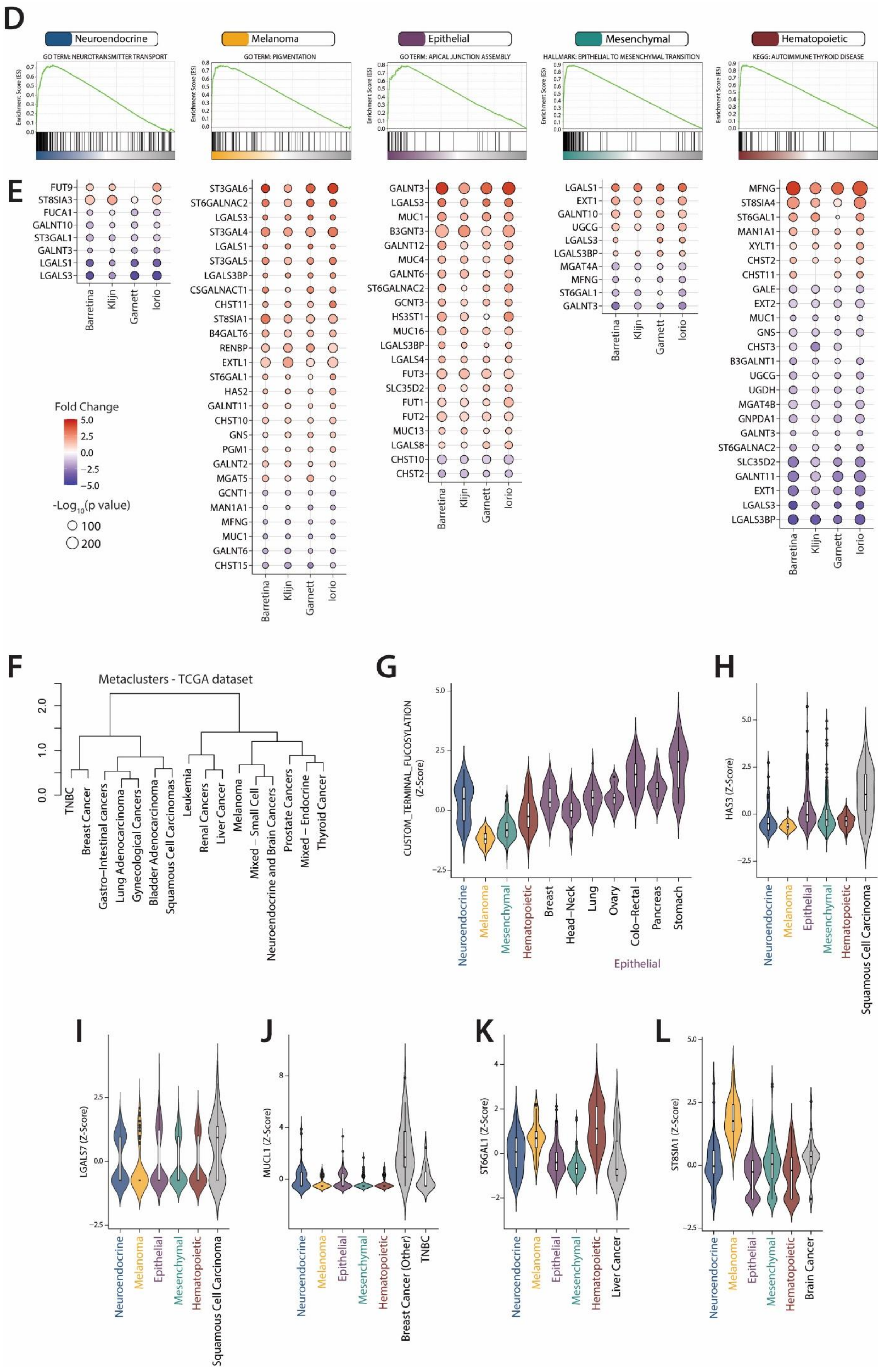
D
Neuroendocrine
GO TERM: NEUROTRANSMITTER TRANSPORT
Melanoma
GO TERM: PIGMENTATION
Epithelial
GO TERM: APICAL JUNCTION ASSEMBLY
Mesenchymal
HALLMARK: EPITHELIAL TO MESENCHYMAL TRANSITION
Hematopoietic
KEGG: AUTOIMMUNE THYROID DISEASE
Enrichment Score (ES)
E
FUT9
ST8SIA3
FUCA1
GALNT10
ST3GAL1
GALNT3
LGALS1
LGALS3
Barretina
Klijn
Garnett
Iorio
ST3GAL6
ST6GALNAC2
LGALS3
ST3GAL4
LGALS1
ST3GAL5
LGALS3BP
CSGALNACT1
CHST11
ST8SIA1
B4GALT6
RENBP
EXTL1
ST6GAL1
HAS2
GALNT11
CHST10
GNS
PGM1
GALNT2
MGAT5
GCNT1
MAN1A1
MFNG
MUC1
GALNT6
CHST15
GALNT3
LGALS3
MUC1
B3GNT3
GALNT12
MUC4
GALNT6
ST6GALNAC2
GCNT3
HS3ST1
MUC16
LGALS3BP
LGALS4
FUT3
SLC35D2
FUT1
FUT2
MUC13
LGALS8
CHST10
CHST2
LGALS1
EXT1
GALNT10
UGCG
LGALS3
LGALS3BP
MGAT4A
MFNG
ST6GAL1
GALNT3
MFNG
ST8SIA4
ST6GAL1
MAN1A1
XYLT1
CHST2
CHST11
GALE
EXT2
MUC1
GNS
CHST3
B3GALNT1
UGCG
UGDH
MGAT4B
GNPDA1
GALNT3
ST6GALNAC2
SLC35D2
GALNT11
EXT1
LGALS3
LGALS3BP
Fold Change
5.0
2.5
0.0
−2.5
−5.0
-Log10(p value)
100
200
F
Metaclusters - TCGA dataset
TNBC
Breast Cancer
Gastro−Intestinal cancers
Lung Adenocarcinoma
Gynecological Cancers
Bladder Adenocarcinoma
Squamous Cell Carcinomas
Leukemia
Renal Cancers
Liver Cancer
Melanoma
Mixed − Small Cell
Neuroendocrine and Brain Cancers
Prostate Cancers
Mixed − Endocrine
Thyroid Cancer
G
CUSTOM_TERMINAL_FUCOSYLATION (Z-Score)
Neuroendocrine
Melanoma
Mesenchymal
Hematopoietic
Breast
Head−Neck
Lung
Ovary
Colo−Rectal
Pancreas
Stomach
Epithelial
H
HAS3 (Z-Score)
Neuroendocrine
Melanoma
Epithelial
Mesenchymal
Hematopoietic
Squamous Cell Carcinoma
I
LGALS7 (Z-Score)
Neuroendocrine
Melanoma
Epithelial
Mesenchymal
Hematopoietic
Squamous Cell Carcinoma
J
MUCL1 (Z-Score)
Neuroendocrine
Melanoma
Epithelial
Mesenchymal
Hematopoietic
Breast Cancer (Other)
TNBC
K
ST6GAL1 (Z-Score)
Neuroendocrine
Melanoma
Epithelial
Mesenchymal
Hematopoietic
Liver Cancer
L
ST8SIA1 (Z-Score)
Neuroendocrine
Melanoma
Epithelial
Mesenchymal
Hematopoietic
Brain Cancer

**Supplementary Figure 9. Characterization of different Glycosylation-related clusters in cancer cell lines.** A) Confusion matrix of the classification of glycosylation clusters in the individual data sets compared with the consensus clusters. The percentage of the same classification between the two clustering are stated at the top of each matrix. B) TSNE plot of cell lines in the different data sets colored by organ of origin and consensus cluster. C) Quantification of consensus cluster associated with organ of origin of the cell lines. D) GSEA plots of selected gene sets enriched in each glycosylation cluster. E) Dot plot of the differential expression of glycosylation-related in glycosylation clusters in each data set. F) Dendrogram displaying the results of the metacluster obtained from the hierarchical clustering of tissue samples. Expression of the geneset “CUSTOM_TERMINAL_FUCOSYLATION” (G) or the genes HAS3 (H), LGALS7 (I), MUCL1 (J), ST6GAL1 (K) and ST8SIA1 (L) in the different subtypes of cell lines. In violin plot: data presented as boxplot indicate the median, 25th and 75th percentiles (hinges) and whiskers represent 1.5 times the interquantile range.

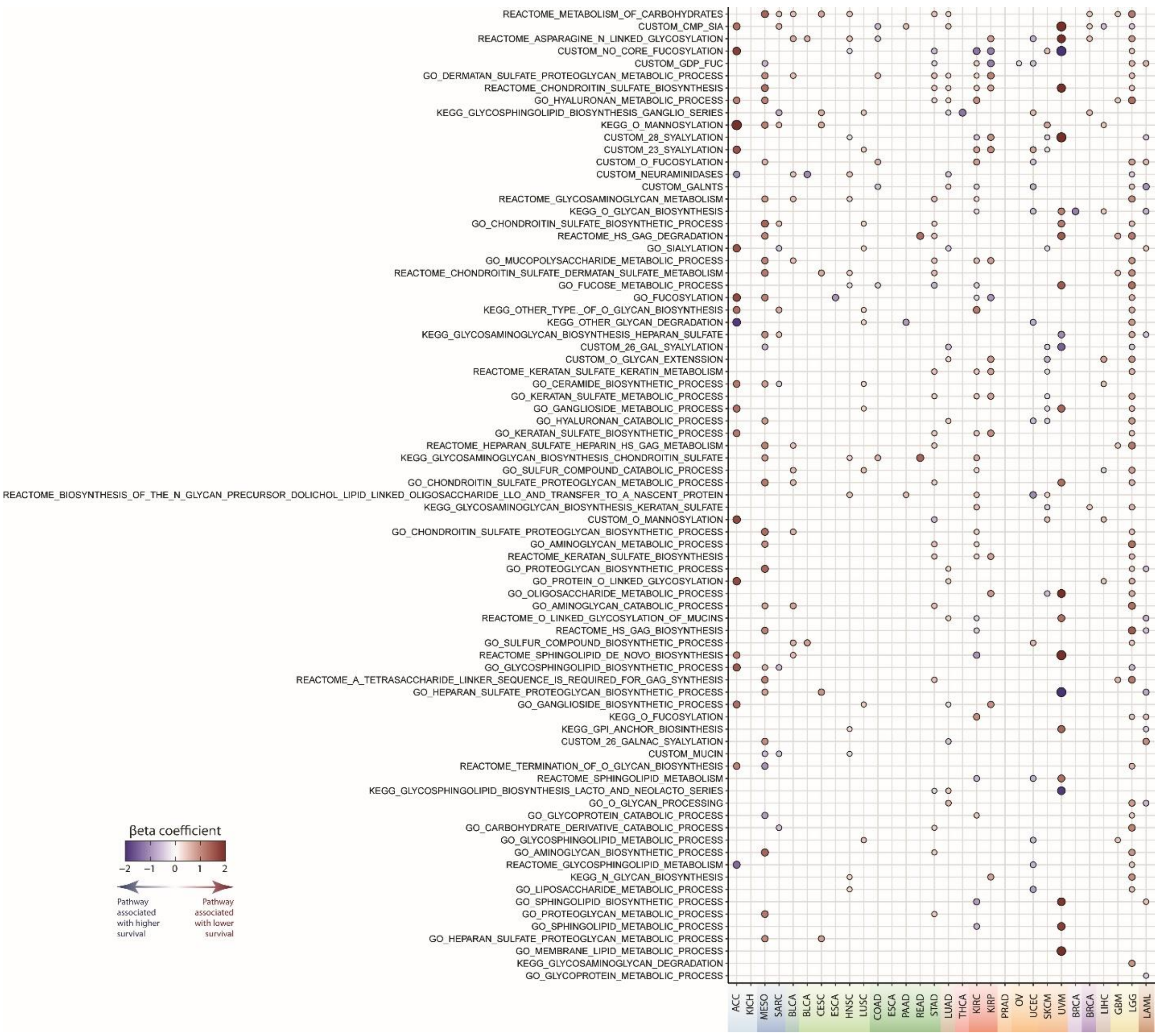

**Supplementary Figure 10. Association of glycosylation pathways with survival in different cancer types.**
